# A C-terminal Processing Protease Implicated in Flagellin Turnover and Developmental Progression in a Bacterial Predator

**DOI:** 10.64898/2026.06.08.730854

**Authors:** Christopher J. Harding, Anali Migueles-Lozano, Carey Lambert, Rob Till, Paul Radford, Kate A. Hadley, Oliver J. Daubney, Anna F. A. Peacock, Anish Parmar, Ishwar Singh, Simon G. Caulton, R Elizabeth Sockett, Andrew L. Lovering, Sampriti Mukherjee

## Abstract

Carboxy-terminal processing proteases (CTPs) are widely conserved bacterial proteases implicated in protein maturation, quality control, and stress responses, yet many family members functions remain unclear. Here, we characterise Bd0967, a previously unstudied CTP from the predatory bacterium *Bdellovibrio bacteriovorus*, and its role in flagellar function during predatory development. Crystal structures reveal a self-compartmentalised protease in which a PDZ domain forms a lid over a large internal cavity accessed through a proteolytic tunnel. Co-purifying peptides occupied two distinct substrate-binding sites, suggesting a coordinated recognition mechanism. Affinity pulldowns coupled with mass spectrometry identified several *Bdellovibrio* flagellins as candidate substrates, which were subsequently validated by biochemical assays and shown to be selectively degraded through recognition of a conserved C-terminal motif. A Bd0967-mCherry translational fusion localised to periplasmic foci during the intracellular predatory growth, consistent with flagellar resorption and flagellin turnover following prey invasion. Deletion of *bd0967* caused developmental defects, including aberrant *Bdellovibrio* cell morphology and reduced predation efficiency. Together, these findings establish Bd0967 as a specialised CTP that couples flagellin degradation and developmental progression in a predatory bacterium.

## Introduction

Carboxy-terminal processing proteases (CTPs) are a widely conserved family of proteolytic enzymes found across bacteria, archaea, and eukaryotes. In bacteria, CTPs are typically located within the periplasm, where their function is to proteolytically process proteins, at or near their C-termini. Although originally identified and characterised in the context of protein maturation and quality control, it has become increasingly clear that CTPs are central to diverse and often crucial cellular processes^1^. Despite their conservation and key physiological functions, most bacterial CTPs remain poorly understood, with their substrate specificity and regulatory mechanisms yet to be uncovered.

Across bacterial species, CTPs have evolved diverse roles, in protein processing, cell signalling and protein quality control. Often these enzymes have highly specialised roles, which have been tailored to lifecycle strategies. For example, in *Bacillus subtilis*, CtpB controls sporulation by cleaving a segment of the C-terminus from the SpoIVFA regulator, triggering the onset of late-stage spore development^2^. Growing evidence now implicates CTPs in stress responses and in critical cell wall maintenance. In *Escherichia coli* the CTP, Prc, when in complex with the outer membrane protein NlpI, maintains cell envelope homeostasis by modulating the proteolytic turnover of peptidoglycan hydrolases, particularly MepS^3^. Similarly, *Pseudomonas aeruginosa* CtpA, which is required for virulence, degrades multiple cell wall hydrolases, mediated via the lipoprotein, LbcA^4^. CTPs also proteolytically regulate stress responses, cell division and host-pathogen interactions in various other bacteria, such as *Staphylococcus aureus*^5^*, Acinetobacter baumannii*^6,7^ *and Chlamydia trachomatis*^8^. Together, these studies highlight that CTPs function not simply as degradative enzymes, but as specialised regulators of bacterial physiology and lifestyles.

Structurally, these CTP enzymes share a common catalytic S41 family protease domain with a serine-lysine catalytic dyad and a regulatory PDZ domain. Despite their shared central architecture, CTPs display striking structural and functional differences, reflecting their wide-ranging physiological roles. The PDZ domain is a key regulatory element, mediating substrate recognition and modulating proteolytic activity^9^. In some enzymes, such as *B. subtilis* CtpB, the PDZ domain functions as an autoinhibitory gate that blocks access to the protease active site, until substrate binding to the PDZ induces its conformational rearrangement, relieving inhibition and permitting specific proteolysis. In contrast, in *E. coli* Prc, substrate engagement triggers a conformational change in the PDZ domain which repositions catalytic residues, in the S41 domain, and activates proteolysis^10^. Additional layers of regulation are provided by interactions with additional adaptor proteins. For instance, *E. coli* Prc forms a stable higher-order complex with the outer membrane lipoprotein NlpI, directing proteolytic activity toward specific cell wall hydrolases at outer membrane locations^11,12^. Similarly, activation of *P. aeruginosa* CtpA requires complex formation with the spiralled outer membrane lipoprotein LbcA^4^, which facilitates periplasmic substrate recognition. LbcA binding induces repositioning of the PDZ domains and coordinates rearrangement of the substrate-binding pocket and catalytic residues across adjacent subunits. These examples underscore how evolution has repurposed the CTP scaffold for distinct regulatory mechanisms and substrate repertoires. The structural differences reflect the intricate and specialised regulatory pathways in which CTPs operate across bacterial species. However, most bacterial CTPs remain uncharacterised, and their broader physiological significance is yet to be uncovered.

In this study, we characterise Bd0967, a previously unstudied carboxyl-terminal processing protease encoded by the predatory bacterium *Bdellovibrio bacteriovorus*, a species with a unique biphasic predatory developmental lifecycle^13^. In liquid media, the lifecycle begins with a highly motile, non-replicative predator that uses its single, membrane-sheathed flagellum to locate and collide with a Gram-negative prey cell. The helical flagellum of *B. bacteriovorus* HD100 is constructed from six distinct flagellin monomer types (Bd0408, Bd0410, Bd0604, Bd0606, Bd3052, and Bd3342) that interact in a spiral via adjacent N- and C-termini to form a semi-rigid, hollow, helical filament with a characteristic dampened wave form, tapering at the tip^14–16^. Apart from Bd0410, whose precise position within the filament remains unresolved, the remaining flagellins occupy defined proximal to distal positions^17^. The filament is rotated by a membrane-embedded motor complex and is ensheathed within a membrane continuous with the bacterium’s outer membrane^18,19^. Upon prey invasion, flagellar rotation ceases and the predator enters the prey periplasm, where it transitions into a sessile replicative form within a dead, rounded prey cell, collectively named a bdelloplast.. Recent cryo-electron tomography studies demonstrated that the flagellum is not shed, as previously proposed, but instead rapidly resorbed into the periplasm of the predator during invasion^20^. This unusual physiological process would require coordinated filament disassembly machinery capable of processing internalised flagellar components. After consuming the prey’s cellular contents and completing replication, the progeny each rebuild a flagellum, regain motility, lyse the prey cell, and resume the free-swimming, non-replicative hunting phase.

Here, affinity-based pulldown assays coupled with mass spectrometry identified multiple *B. bacteriovorus* flagellins as Bd0967-interacting proteins, which were subsequently validated as direct substrates using proteolytic degradation assays. Structural characterisation of Bd0967, reveals a divergent CTP architecture containing an extended scaffold and large internal cavity. Mechanistically, we demonstrate that Bd0967 selectively recognises and degrades monomeric flagellins via their C-terminal regions in vitro. In addition, deletion of Δ*bd0967* resulted in aberrant bdelloplasts, and impaired predation efficiency. Furthermore, a Bd0967-mCherry fusion localises to discrete puncta, inside *B. bacteriovorus*, during the intracellular predatory growth phase. These findings establish a direct link between CTP-mediated proteolysis, by Bd0967, and flagellar-filament disassembly, which is a critical developmental transitionary event required for the shift to intracellular growth by the predator. Our findings uncover a previously unknown function for bacterial CTPs in structural remodelling and lifecycle control, expanding the functional diversity of this important protease family.

## Results

### Bd0967 structure exhibits hallmarks of CTP protease

We determined the structure of WT Bd0967, from *B. bacteriovorus* HD100, as well as the catalytically inactive Bd0967_S457A_, to 2.04 Å and 2.58 Å, respectively. The electron density maps were well-resolved from residues 29 to 665, encompassing the full secreted form of Bd0967, with only minor regions of disorder. Bd0967 crystallised as a monomer, with four molecules present in the asymmetric unit (ASU), in P1 space group. Each monomer adopts a nearly identical bowl-like conformation featuring a hinged lid, together forming a large, self-enclosed chamber.

Bd0967 is composed of four distinct domains: an N-terminal domain (NTD), a C-terminal domain (CTD), a protease domain, and a PDZ domain (Figure 1A). The NTD and CTD are positioned at opposite ends of the central protease core and extend outward, forming the scaffold of the bowl-shaped architecture. This scaffold is stabilised by a composite β-sheet formed by a single β-strand contributed by both the NTD and CTD at the distal end of the α-helices and two individual β-strands. The CTD, consists of five α-helices and a single β-strand, and adopts an open paddle-like conformation that is structurally distinct from CTDs found in other C-terminal processing proteases.

**Figure 1.**
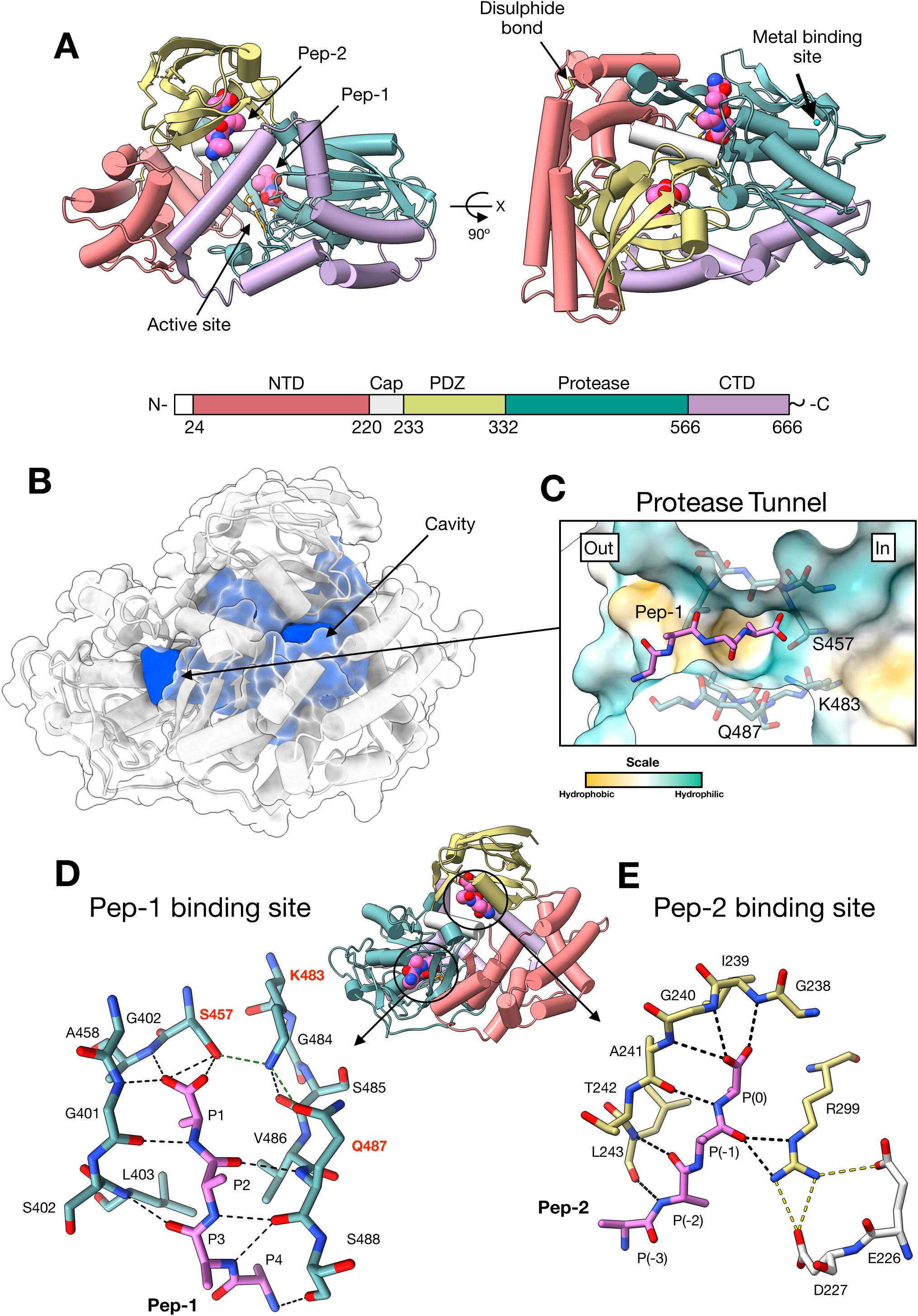
Structure of the Bd0967 self-compartmentalised protease. **(A)** Overall crystal structure of Bd0967. The four structural domains are colour coded according to the schematic. There is an N-terminal domain (NTD), central protease domain, C-terminal domain (CTD), and a PDZ domain. The three core domains form a bowl-shaped scaffold that is capped by the PDZ domain, generating a self-compartmentalised proteolytic chamber. Active site residues highlighted in orange. **(B)** Internal cavity (blue), calculated with KVFinder, revealing the extensive enclosed chamber and connecting protease tunnel. **(C)** The ∼15 Å protease tunnel formed by the peptide-binding channel of the protease domain which is formed between the Cap subdomain and protease domain. The tunnel contains multiple hydrophobic pockets for substrate selection. The catalytic residues, S457 & K483 are located at the end of the tunnel. The co-purified peptide (pink), modelled as a poly alanine chain, is shown within the tunnel. **(D)** Bd0967 peptide binding sites are highlighted on the structure, which is orientated in the identical pose as for the surface illustration in panel B. Illustration of the hydrogen-bonding network at the Pep-1 binding site. Residues from the protease domain (blue), which interact with the bound peptide (pink), are shown. Catalytic residues are highlighted by red text. Black dashed lines represent H-bonds and green dashed lines show catalytic residue H-bonds. **(E)** Shows the hydrogen-bonding network at the Pep-2 binding site, with PDZ residues (yellow), Cap residues (silver), and bound peptide (pink). R299, from the PDZ domain, mediates key interactions with Pep-2 and the Cap subdomain. Dashed lines represent H-bonds.

The protease domain adopts a characteristic fold of S41 family proteases, in which two mixed beta sheet regions are separated by a series of α-helices, forming a channel with the catalytic residues located at the base. A single α-helix acts as an independent subdomain (Cap), which links the NTD and PDZ domains.

The overall architecture is completed by a PDZ domain, composed of a curved five-stranded anti-parallel β-sheet, flanked by two α-helices. This domain is connected to the protease domain and Cap subdomain, at a single point, via a flexible hinge and sits above the bowl-shaped scaffold, acting as a lid that encloses a large internal cavity (Figure 1B). This cavity is a defining structural feature of Bd0967 and has not been described in other CTPs. Using KVFinder^21^, the cavity volume was estimated at 13952 Å^3^ with a surface area of 3979 Å^2^. The cavity is approximately spherical, with a diameter of ∼30 Å, which could theoretically accommodate small, unfolded proteins, or regions of larger substrate, within a protected environment.

Positioned beneath the PDZ lid, the Cap subdomain sits directly above the catalytic site and converts the protease channel into a tunnel (Figure 1C). This arrangement provides a narrow route by which substrates must pass through into the enclosed cavity, passing over the catalytic centre, as they do so.

As in other CTPs, Bd0967 contains a catalytic dyad, in which S457 acts as the nucleophile and K483 functions as a general base, activating S457 and the water molecule required for collapse of the tetrahedral intermediate^22^. K483 also forms a hydrogen bond with N487, in addition to interactions with the backbone carbonyl of S485, stabilising the orientation and charge during catalysis.

In addition, Bd0967 contains two structural features not observed in other characterised CTPs. First, it harbours a metal-binding site within a peripheral loop of the protease domain (Figure 1A & SI Figure 1A), where a metal ion is coordinated by seven oxygen atoms: one from a water molecule, the side-chain carboxyl of D436, and backbone carbonyls from G411, F414, Q415, D438, and M440. Residue P439 appears to position these carbonyls to facilitate coordination. Based on coordination geometry^23^, bond length, valence and electron density, this metal ion is most likely calcium. This site likely serves a structural role in the protease domain, by stabilising an otherwise flexible loop region.

Second, Bd0967 features a disulfide bond within the NTD, linking C33 and C100 (SI Figure 1B). C33 lies at the N-terminus and the disulfide bond likely stabilises this region and, by extension, the overall scaffold of the NTD. The presence of a bona fide disulfide bond, coupled with predicted signal peptide (SI Figure 2A), supports localisation to the predator’s periplasm, where the enzyme would have access to substrate proteins.

The distribution of B-factor, flexibility values across Bd0967 varies significantly between domains (SI Figure 3A). Lower B-factors are observed in the protease domain and adjacent regions, indicating relative rigidity. In contrast, higher B-factor values are found in the distal portions of the NTD and CTD, as well as in the PDZ domain, suggesting these regions are more flexible and may undergo greater conformational sampling. This flexibility supports the idea that the PDZ domain may adopt multiple conformations as part of regulating the catalytic mechanism, which has been observed previously for enzymes in this protease family (SI Figure 4). We observed subtle conformational differences in the PDZ domain between different Bd0967 molecules comprising the asymmetric unit and between the WT and the catalytically inactivated S457A mutant structures. These conformational differences corresponded to an approximately ∼8° rotation about the hinge region (SI Figure 3). In all cases, the PDZ domain adopted a peptide-bound, open form. While this shift is modest compared to the larger PDZ domain movements reported in other CTPs, it highlights the inherent flexibility of the domain and suggests that additional conformations may be sampled but not captured in our structures.

### Bd0967 has two distinct peptide binding sites

Bd0967 has two distinct peptide binding sites: site 1 within the protease tunnel of the S41 domain, formed between the protease domain and Cap subdomain; and site 2 within the PDZ domain (Figure 1A). Both sites exhibit clear electron density corresponding to co-purified and co-crystallised substrate peptides, which are referred to as Pep-1 and Pep-2, respectively (SI Figure 1C & D). These peptides most likely originate from *E.coli* proteins, present during heterologous expression of Bd0967.

In both WT Bd0967 and Bd0967_S457A_ structures, this substrate peptide density is consistent with approximately four amino acid residues and is comparable in quality to the main-chain density of the protein, allowing confident modelling of the substrate peptide backbone. However, side-chain density was not well resolved, likely reflecting a combination of moderate resolution and possible heterogeneity in peptide identity across protomers. Accordingly, the peptides were conservatively modelled as poly-alanine, although residual positive F_o_-F_c_ density suggests the presence of larger amino-acid side chains. We addressed this later by synthetic peptide binding experiments described below.

Despite efforts to identify the peptide by mass spectrometry, the sequence of the interacting peptides, from the heterologous *E. coli* proteome, could not be determined. Notably, the peptide density is highly similar in both WT and catalytically inactive (S457A) structures, which adopt comparable “open” PDZ conformations. At the current resolution, it was not possible to definitively distinguish between a continuous peptide substrate and a cleavage product. However, the presence of well-defined electron density consistent with a C-terminal carboxylate in the WT structure, located proximal to the catalytic serine, contrasts with its absence in the Bd0967_S457A_ structure. This suggests that the WT structure likely contains a cleaved peptide product, whereas the mutant retains a continuous peptide, without the cut carboxylate site.

Together, these observations indicate that both binding sites support stable, ordered interactions with peptide backbones spanning approximately 4 amino acids. Beyond the core substrate binding pockets, electron density became poorly resolved, suggesting the flanking peptide regions are either more variable and/or become increasingly disordered (adopting multiple conformations).

The protease-bound co-purified peptide (Pep-1) makes extensive hydrogen bonding contacts with backbone atoms of the Bd0967 peptide-binding tunnel. Pep-1 is positioned between two β-strands, creating a short, three-residue augmented β-sheet interaction (Figure 1D). The S1 pocket, which accommodates the P1 residue of the bound peptide, is formed by the nonpolar residues L403, I461, V486, and F506, and likely favours small hydrophobic side chains at this position. In contrast, the S2 pocket, formed by F220, F228, M232, V489, and K500, is broader and more solvent-accessible, suggesting that it can accommodate larger side chains at the P2 position. The P3 and P4 positions are more exposed to solvent beyond the tunnel entrance, making them more permissive to polar residues.

The co-purified peptide bound to at the PDZ-domain (Pep-2), binds to a relatively hydrophobic pocket on the internal surface of the PDZ domain. Its C-terminus carboxyl group is anchored at the back of the pocket by the carboxylate-binding loop (G238-I239-G240-A241), as observed in other PDZ-peptide complexes^2,10^. The peptide forms an extended β-sheet interaction with β-strand 4 of the PDZ domain, making hydrogen bond contacts with the backbone atoms of I239, G240, A241, T242 and L243. The sidechain of R299 runs parallel to the peptide chain, mimicking a β-strand interaction and stabilising the P0 and P-1 positions. Notably, R299 sidechain occupies a key structural position, forming both a salt bridge and hydrogen bonds with E226 and D227 of the protease cap (Figure 1E). These contacts likely contribute to allosteric communication between peptide binding and conformational state, consistent with mechanisms proposed for other CTP family members^2^.

### Bd0967 associates with *B. bacteriovorus* flagellins

To this point, all peptide interactions observed for Bd0967 involved co-purified peptides originating from the *E. coli* expression host. To identify potential substrates and native proteins which interact with Bd0967 we performed a pulldown assay using immobilised Bd0967_S457A_ and fractionated *Bdellovibrio bacteriovorus* HD100 predator cell lysates, comprising a soluble fraction enriched in cytoplasmic and periplasmic proteins, and an insoluble fraction enriched in membrane-associated proteins. Samples were evaluated by SDS-PAGE, alongside controls, to identify presence of Bd0967-interacting proteins (Figure 2A). Unique protein bands, corresponding to proteins with masses ranging between 20-35 KDa, were observed when Bd0967_S457A_ was incubated with *B.b*.HD100 lysate insoluble membrane fraction (S1 = Bd0967-bound beads + HD100 lysate). These bands were not visible in controls (C1 = unbound beads + HD100 lysate & C2 = Bd0967-bound beads + lysis buffer), or in the Bd0967-bound beads + HD100 soluble material lysate sample (Figure SI 5).

**Figure 2.**
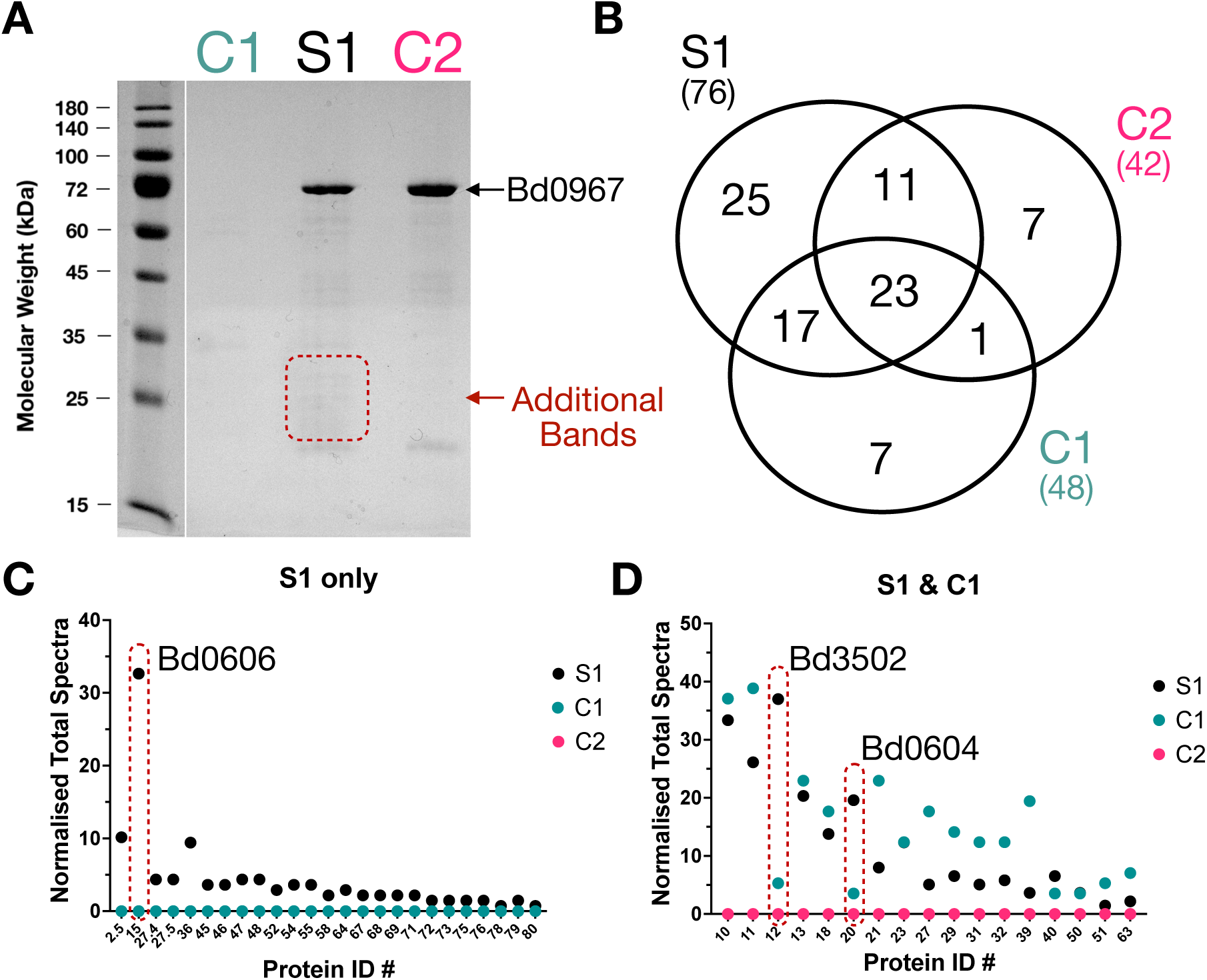
Pulldown assays. **(A)** SDS-PAGE of pulldown with Bd0967_S457A_ as bait protein with *Bdellovibrio bacteriovorus* HD100 lysate (insoluble fraction). S1 = Bd0967-bound beads + HD100 lysate, C1 = unbound beads + HD100 lysate & C2 = Bd0967-bound beads + lysis buffer. Additional bands are seen between the 20-35 kDa range. **(B)** Venn diagram illustrating the Unique proteins identified in each in-gel trypsin digested sample analysed by mass spectrometry. Numbers in brackets refers to the total number of proteins detected for each sample. **(C)** Graph showing the normalised total spectra, for unique samples for S1. Bd0606 (highlighted by the red dashed shape) had the highest normalised spectra. **(D)** Graph showing the normalised total spectra, for samples identified in S1 & C1. The flagellins Bd3502 & Bd0604 (highlighted by the red dashed shapes) were highly enriched in the S1 samples and detected at low levels in the C1 sample.

To identify these proteins, an in-gel trypsin digest of proteins contained in the whole lanes, followed by liquid chromatography mass spectrometry (LC-MS), was performed to identify the Bd0967-interacting protein species. LC-MS results were analysed by Scaffold^24^, which identified a total of 91 unique protein spectra across the three samples S1, C1 & C2. Of these 91 protein spectra, 25 were unique to S1, based on normalised total spectra counts (Figure 2B, Supporting Document 1). We disregarded contaminating proteins originating from *E. coli* or human sources, besides proteins predicted to be localised to the cytoplasm, as these would be unlikely to encounter Bd0967, predicted to be localised to the periplasm based on its signal peptide and bonafide disulphide linkage. This left 7 candidates, of which a flagellin (Bd0606) showed the highest enrichment, with a high normalised total spectra count of 33, whereas other candidate proteins in this group, had substantially lower spectra counts (<10) (Figure 2C). Additionally, two other flagellins (Bd0604 and Bd3052) were also enriched in S1 and detected at low levels in the C1 control (Figure 2D). These three proteins (Bd0604, Bd0606 & Bd3052) all belong to the same flagellar filament apparatus^15,17^ and possess similar C-termini sequences (MKLV-, LRLIG & LKLIG, respectively), suggesting a tangible link between Bd0967 and flagellar components as a proteolytic target. The presence of flagellins being present in the insoluble membrane fraction is consistent with the *B. bacteriovorus* flagellum being external to the cytoplasmic membrane and outer-membrane sheathed.

### Bd0967 degrades the flagellin Bd0606 *in vitro*

To validate Bd0606 as a substrate of Bd0967, we performed in vitro degradation assays using purified Bd0606 monomers. WT Bd0967 efficiently degraded Bd0606, under standard reaction conditions, whereas a catalytic null-mutant Bd0967_S457A_, showed no activity (SI Figure 6), confirming degradation is dependent on catalytic activity.

Time-course analysis revealed the progressive formation of discrete intermediate fragments prior to their eventual disappearance (Figure 3A). The transient accumulation of these intermediates suggests that Bd0967 does not degrade the substrate in a single continuous event like Clp proteases^25^. Instead, it generates cleavage products that are subsequently processed further. This behaviour is consistent with progressive substrate cleavage involving release and subsequent re-engagement of intermediate products.

**Figure 3.**
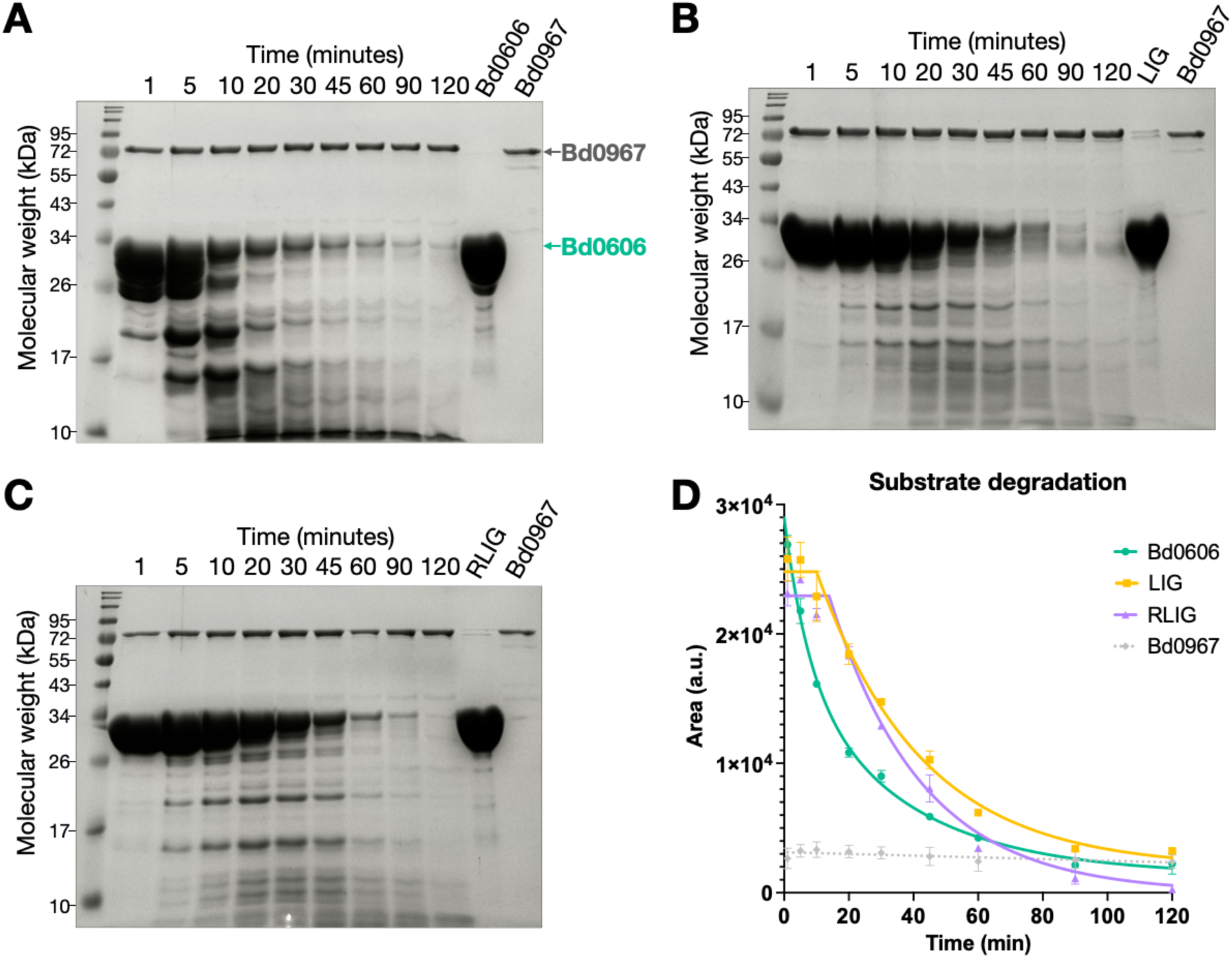
Bd0967 degrades the flagellin Bd0606. **(A)** Proteolytic activity assay monitoring the degradation of substrate, Bd0606, over time. Bd0967 rapidly degrades Bd0606 under standard assay conditions. Distinct lower molecular weight intermediates are formed. Bd0967 and Bd0606 species are marked in the gel for clarity. **(B)** Proteolytic activity assay monitoring the degradation of substrate, Bd0606-LIG, over time. **(C)** Proteolytic activity assay monitoring the degradation of substrate, Bd0606-RLIG, over time. **(D)** Plot of gel band intensity overtime for each Bd0606 species. Bd0606 is the more efficient substrate that does not have a plateau phase to its degradation. Experiments were performed in triplicate, showing mean and SD error.

Notably, the proteolytic activity of Bd0967 towards Bd0606 appears substantially greater than that reported for previously characterised CTPs, many of which exhibit relatively slow turnover rates, particularly in the absence of adaptor / activator proteins. Notably, Bd0967 displays robust activity without the requirement for accessory activation factors. To the best of our knowledge, this study also represents the first reported CTP degradation assay performed under conditions of large substrate excess relative to enzyme (100:1 molar ratio). In contrast, previous studies of related systems employed considerably lower substrate-to-enzyme ratios, including 14:1 for MepS:Prc^10^ and approximately 1:1 for PA1198:CtpA^4^.

Despite this substantial substrate excess, Bd0967 efficiently degrades Bd0606 to completion in comparable time periods, highlighting its unusually high proteolytic capacity. This elevated activity could be advantageous for rapid turnover of abundant substrates such as resorbed flagellins following filament retraction into the periplasm^20^.

### C-terminal features of flagellins contribute to substrate recognition

Sequence comparison of all six *B. bacteriovorus* flagellins revealed substantial conservation across the full-length proteins (50–70% sequence identity). Notably, this conservation extends to the C-terminal region, which is expected to interact with the Bd0967 PDZ domain. The terminal residues exhibit a highly conserved register, following the motif [LM]-[KR]-L-[ILV]-G (SI Figure 9B). Bd0408, the most proximal flagellin, is a notable exception, having an extended C-terminal region of six additional residues that deviates from this conserved motif.

To directly test the contribution of the C-terminus, we generated truncation mutants of Bd0606 lacking the final three or four residues, referred to hereafter as ΔLIG and ΔRLIG, respectively. Surprisingly, WT Bd0967 retained an ability to degrade both variants. However, turnover efficiency of ΔLIG and ΔRLIG was markedly reduced in comparison to WT Bd0606 (Figure 3B & 3C). Both ΔLIG and ΔRLIG appear to have a stalled phase where turnover is limited. This stalled phase lasts for approximately 20 minutes under standard reaction conditions, after which, substrate turnover is rapid and efficient. The difference in turnover efficiency is illustrated in Figure 3D, where the peak area of Bd0606 bands were plotted against time. WT Bd0606 is degraded exponentially, whereas ΔLIG and ΔRLIG turnover has a distinct plateau followed by exponential decay. In addition, the intermediates observed on the SDS-PAGE, accumulate more quickly and in more abundance for WT Bd0606 in comparison to ΔLIG and ΔRLIG truncation mutants.

To further probe the substrate recognition mechanism, we synthesised two peptides: NAMPNSALRLIG (Inhibitor-1), corresponding to the C-terminus residues of Bd0606; and NAMPNSALRIG (Inhibitor-2), a slight variant lacking the third residue (L) from the C-terminal end. These peptides mimic the C-terminal sequence of the Bd0606 substrate, so they were hypothesised to bind directly with the PDZ domain, thus acting as competitive inhibitors. Using the same gel-based assay under standard reaction conditions, we examined the effect of increasing concentration of peptide on the activity of Bd0967. Indeed, at higher concentrations of Inhibitor-1 (>6mM) the turnover of Bd0606 was drastically reduced (Figure 4A). This suggests that Inhibitor-1 (peptide mimicking WT Bd0606 flagellin C-terminus) acts as a competitive inhibitor, reducing the activity of Bd0967, likely by interacting with the PDZ domain and blocking binding of substrates. Interestingly, Inhibitor-2, which differs by missing a single residue at the C -3 position (no leucine), compared to Inhibitor-1, showed very little inhibitory activity towards Bd0967 (Figure 4B), suggesting correct sequence register is important for binding. The differences in inhibition are illustrated in Figure 4C, through gel band analysis. Together with the reduced turnover efficiency of ΔLIG and ΔRLIG, these findings highlight the importance of the interaction between the C-terminus and the PDZ domain in the recognition and activation of Bd0967, although there appears to be a degree of tolerance.

**Figure 4.**
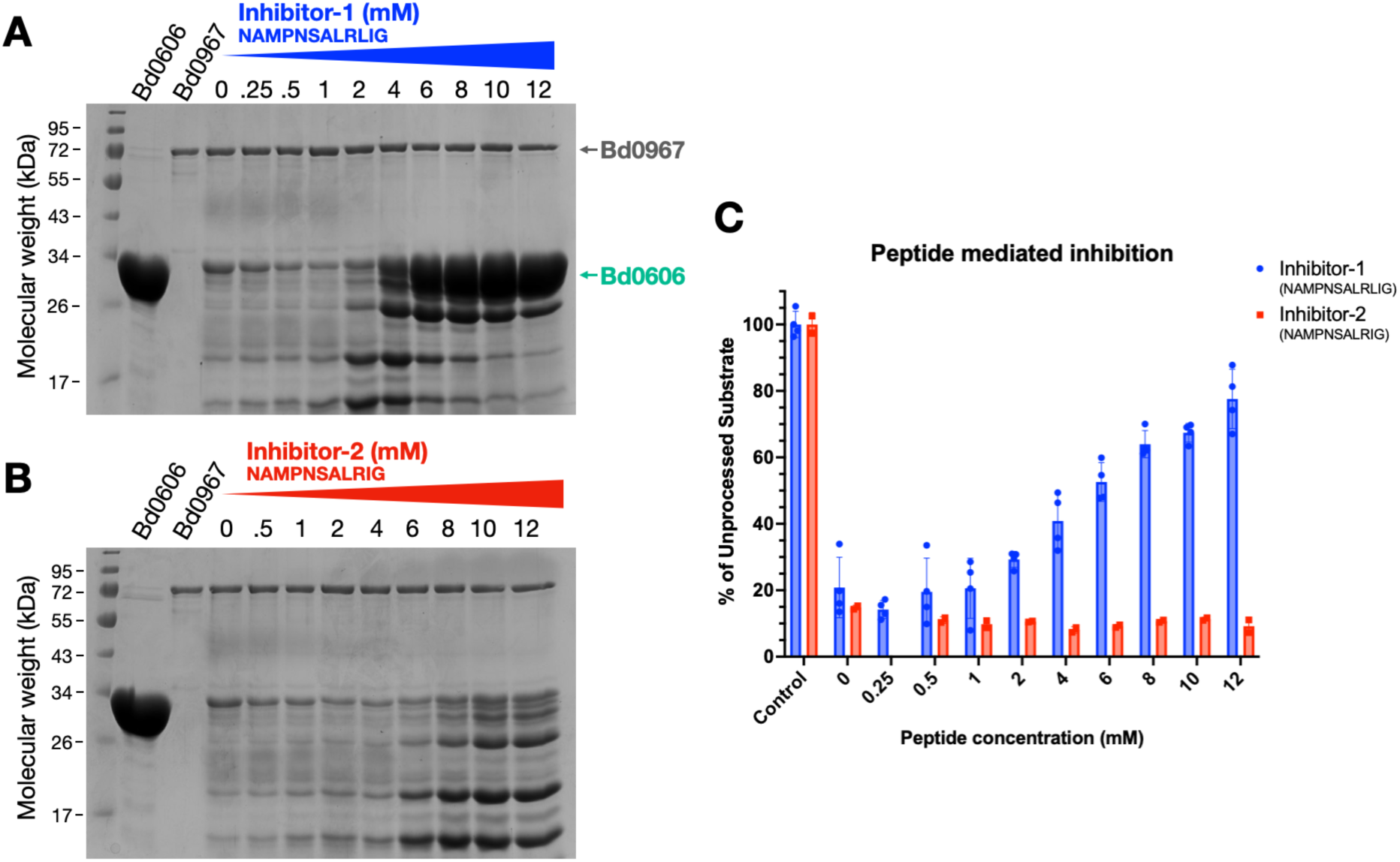
Inhibition of Bd0967 activity by C-term derived peptides. **(A)** SDS-PAGE gel showing the affect of increased peptide concentration Inhibitor-1 (NAMPNSALRLIG, blue - derived from C-terminus of Bd0606), under standard proteolytic assay conditions. Inhibitor-1 acted as a competitive inhibitor preventing degradation of Bd0606 at higher concentrations. **(B)** SDS-PAGE gel showing the affect of increased Inhibitor-2 (NAMPNSALRIG, red, lacking L at the C-3 position), under standard proteolytic assay conditions. Inhibitor-2 does not prevent degradation of Bd0606 even at higher concentrations. **(C)** Plot of gel band intensities (peak area) vs concentration of peptide, illustrating peptide mediated inhibition. Experiments were performed in triplicate, showing mean and SD error.

Corroborating this, an impaired PDZ-domain substrate recognition mutant, Bd0967_R299M_, was incapable of turning over ΔLIG and ΔRLIG substrates and showed limited activity towards WT Bd0606 (SI Figure S6). This finding suggests C-terminal recognition of flagellin substrate by the PDZ domain is critical to induce proteolysis. This critical feature renders the Bd0967_R299M_ mutant very inefficient, which is compounded when tested with an inefficient substrate such as ΔLIG and ΔRLIG.

Together, these results demonstrate that C-terminal interactions contribute significantly to efficient substrate turnover by Bd0967.

### Bd0967 degrades multiple flagellins with differing efficiencies

Having established Bd0606 as a substrate and identified key determinants of recognition, we next assessed whether Bd0967 exhibits activity towards additional diverse flagellins which comprise the flagella filament. The filament is composed of six flagellins arranged from proximal to distal positions^17^ (Figure 5A). From this set, we successfully expressed and purified three additional flagellins: Bd0408 (proximal), Bd0604 (central), and Bd3342 (distal), creating a panel of flagellins which span the full length of the filament. Due to technical limitations of expression and purification, we were unable to obtain the full library of Bdellovibrio flagellins.

**Figure 5.**
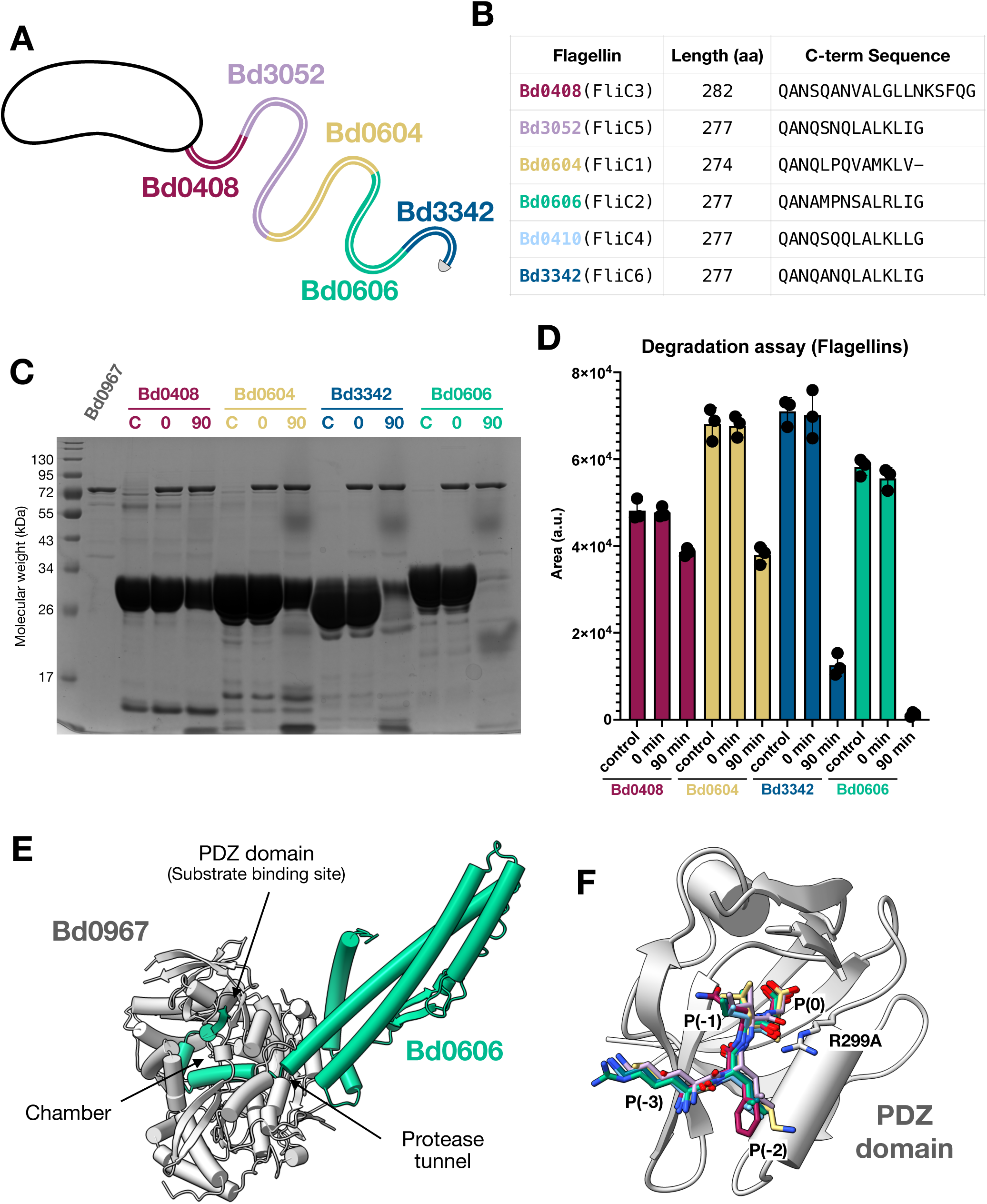
Bd0967 degrades multiple flagellins with differing efficiencies and shared C-terminal recognition features. **(A)** Cartoon representation of flagellar filament architecture illustrating relative location of each flagellin within the filament (adapted from Lida et al). **(B)** Table showing the length and aligned C-terminal sequence of flagellins. Flagellins are colour matched to the cartoon and ordered from proximal to distal. **(C)** SDS-PAGE showing degradative activity of Bd0967 towards different purified recombinant *B. bacteriovours* flagellins. Lanes show a flagellin only control sample. **(C)**, alongside samples taken at 0 and 90 minutes time intervals. Representative image of three biological replicates. **(D)** Chart of band intensities (peak area) for each flagellin protein. Three individual measurements for each point are displayed with error bars displaying corresponding standard deviation. **(E)** AlphaFold3 prediction, modelling the interaction between Bd0967 (silver) and Bd0606 (turquoise). Bd0606 engages the Bd0967 protease domain primarily via helical contacts, while the C-terminal region threads through the protease tunnel into the central cavity, where the C-terminus interacts with the PDZ domain. Comparable interaction predictions between Bd0967 and the other Bdellovibrio flagellins was also observed SI figure. **(F)** The predicted binding mode of the flagellin C-terminus closely mirrors the interaction observed for the co-purified peptide. Although Bd0408 possesses a divergent C-terminus, it is still predicted to bind the PDZ domain, outlining the algorithm’s tendency to match PDZ domains with C-terminal peptide motifs.

Under standard reaction conditions, Bd0967 was capable of degrading all three additional flagellins tested, albeit with differing efficiencies in the 60-minute end point assay (Figure 5C & D). Bd0408 was degraded to a lesser extent compared to the other substrates, whereas Bd0604 and Bd3342 showed more substantial turnover. Although, none of these flagellins were degraded to the same extent as Bd0606 within this timeframe, indicating that Bd0606 represents a more efficient substrate for Bd0967.

To ensure that differences in degradation were not attributable to variations in flagellin monomer protein folding, circular dichroism (CD) spectroscopy was performed. All flagellins displayed similar spectra characteristic of α-helical proteins, with minima at ∼208 nm and ∼222 nm (SI Figure 7). Conversion to mean residue ellipticity (MRE) enabled direct comparison, confirming that the proteins adopt comparable folded states.

Together, these results demonstrate that Bd0967 is capable of degrading multiple *B. bacteriovorus* flagellins, although with varying efficiencies. This expands the substrate repertoire of Bd0967 and suggests that substrate recognition is not limited to a single flagellin, but may instead reflect shared structural features and conserved C-terminal motifs.

### Bd0967 exhibits substrate selectivity

To determine whether Bd0967 acts as a general protease or is selective, we assessed the proteolytic activity of Bd0967 against a limited but diverse panel of non flagellar *Bdellovibrio*-derived proteins, purified in-house from unrelated projects. These proteins were selected to cover variation cellular locations, a range of sizes, sequences and folds. For example, Bd0108 is known to be a small intrinsically disordered protein, found in the periplasm. Under standard assay conditions, none of these proteins were substrates of Bd0967 (SI Figure S8). To determine whether Bd0967 may cleave unfolded or non-native proteins, we also tested proteolytic activity towards native and denatured Hen egg white lysozyme and Bovine serum albumin (BSA), and we observed no cleavage for either form (SI Figure 9A). These results suggest that Bd0967 does not act as a general degradative protease, targeting denatured, or unstructured proteins but instead exhibits selectivity towards its substrate. Also, due to its predicted location in the periplasm, Bd0967 would be limited to periplasmic proteins.

To assess whether the C-terminal motif observed in flagellins is unique within the *Bdellovibrio bacteriovorus* HD100 proteome, we performed a motif-based search of the genome using a permissive consensus sequence, [LMIV]-[RKHMG]-[LIV]-[ILVMFGA]-[GAN]->. This search returned a limited number of candidate proteins (n = 7), including five of the annotated flagellins. The remaining hits corresponded to predicted cytoplasmic or hypothetical proteins with no clear functional relationship to flagellins. These results indicate that the identified C-terminal motif is relatively rare within the HD100 proteome and is strongly associated with flagellins. This supports the idea that Bd0967 recognises a conserved C-terminal feature characteristic of flagellins.

To further explore the molecular basis of Bd0967 substrate recognition, we used AlphaFold3 to model interactions between Bd0967 and each of the six *B. bacteriovorus* flagellins (Bd0408, Bd0410, Bd0604, Bd0606, Bd3052, and Bd3342). All flagellins share similar overall lengths, predicted folds, and C-terminal sequences (Figure 5B).

The models predict a highly conserved mode of interaction across all flagellins (SI Figure 11). In each case, the flagellin engages the Bd0967 protease domain primarily through lower confidence helical contacts (SI Figure 11), while the C-terminal region threaded through the protease tunnel into the central cavity. Within this cavity, the C-terminus is predicted to interact directly with the PDZ domain substrate-binding pocket (Figure 5E).

AlphaFold3 consistently predicts this C-terminal flagellin-PDZ domain engagement with high confidence across all flagellins (Figure 5F), supporting a conserved mode of recognition. The models further suggest that approximately 30–35 residues of the C-terminal region extend into the proteolytic chambre, placing a potential cleavage site near the sequence VAAATA of Bd0606. This region partially aligns with the consensus motif identified for flagellin C-termini, and is consistent with the proposed PDZ-mediated recognition mechanism.

As a specificity control, additional Alphafold2 predictions were performed using Bd0967 and the same panel of proteins, which were not degraded in our biochemical assays (SI Figure 8). None of these models reproduced the characteristic interaction observed with the flagellins, in which the substrate C-terminal region extends through the protease tunnel and engages the PDZ domain. Conversely, all six B. bacteriovorus flagellins were consistently predicted to adopt this binding mode. To further assess the specificity of this interaction, we modelled complexes between the flagellin Bd0606 and the three other predicted PDZ-containing C-terminal processing proteases encoded by *B. bacteriovorus* (Bd0169, Bd1239 and Bd3534). In contrast to Bd0967, none of these proteases were predicted to engage the Bd0606 C-terminus within their PDZ domains or position the substrate for entry into the protease channel. Together, these predictions support a specific interaction between Bd0967 and the Bdellovibrio flagellins that is not readily recapitulated with unrelated proteins or related CTP family members.

### Bd0967 is required for efficient developmental progression in the predatory lifecycle of *B. bacteriovorus*

Bd0967 is predicted to be periplasmic based on its signal peptide and disulfide bond formation (SI Figure 2A). As a first step to characterizing its role in *B. bacteriovorus* HD100, we investigated the spatiotemporal dynamics of Bd0967 during predation. We generated a translational Bd0967-mCherry C-terminal fusion and visualised its localisation by confocal fluorescence microscopy across defined stages of the *B. bacteriovorus* predatory lifecycle. Attack phase (AP) cells were counterstained with the membrane dye Vibrant DiO to delineate membranous cell boundaries and visualize the sheathed flagellum, and predation assays for obtaining growth phase (GP) cells used *mTurquoise2*-expressing *E. coli* prey to identify bdelloplasts by backlit fluorescence microscopy^26^. In free-swimming AP cells, Bd0967-mCherry displayed a non-uniform distribution: approximately 10% of cells exhibited a discrete polar focus, ∼60% displayed a distributed pattern throughout the cell body, with no obvious enrichment at the flagellar pole or elsewhere, and the remainder of cells had undetectable fluorescence (Figure 6A-C). We note that this method is not able to definitively determine the compartmentalisation of Bd0967 due to its initial synthesis in the cytoplasm before transport to a periplasmic location. Upon invasion and after two-hours post bdelloplast formation, Bd0967-mCherry underwent a marked redistribution, with ∼60% of GP cells within bdelloplasts displaying a punctate pattern rather than being distributed throughout the cell body (Figure 6D-E). Because the rounded bdelloplast and the coiled intraperiplasmic predator lack a defined morphological reference axis, we did not attempt to assign puncta to specific subcellular positions within growth-phase cells. Nevertheless, this transition from a partially polarised distribution in attack phase to a discrete punctate pattern during growth phase coincides temporally with the period of flagellar resorption observed by cryo-ET^20^ and is consistent with a model in which Bd0967 is synthesised and transitions from a poised, resting state, in flagellate, motile AP *Bdellovibrio*, to an active configuration upon internalisation of flagellin substrates into the periplasm following filament disassembly and the onset of predatory growth phase by *Bdellovibrio*.

**Figure 6.**
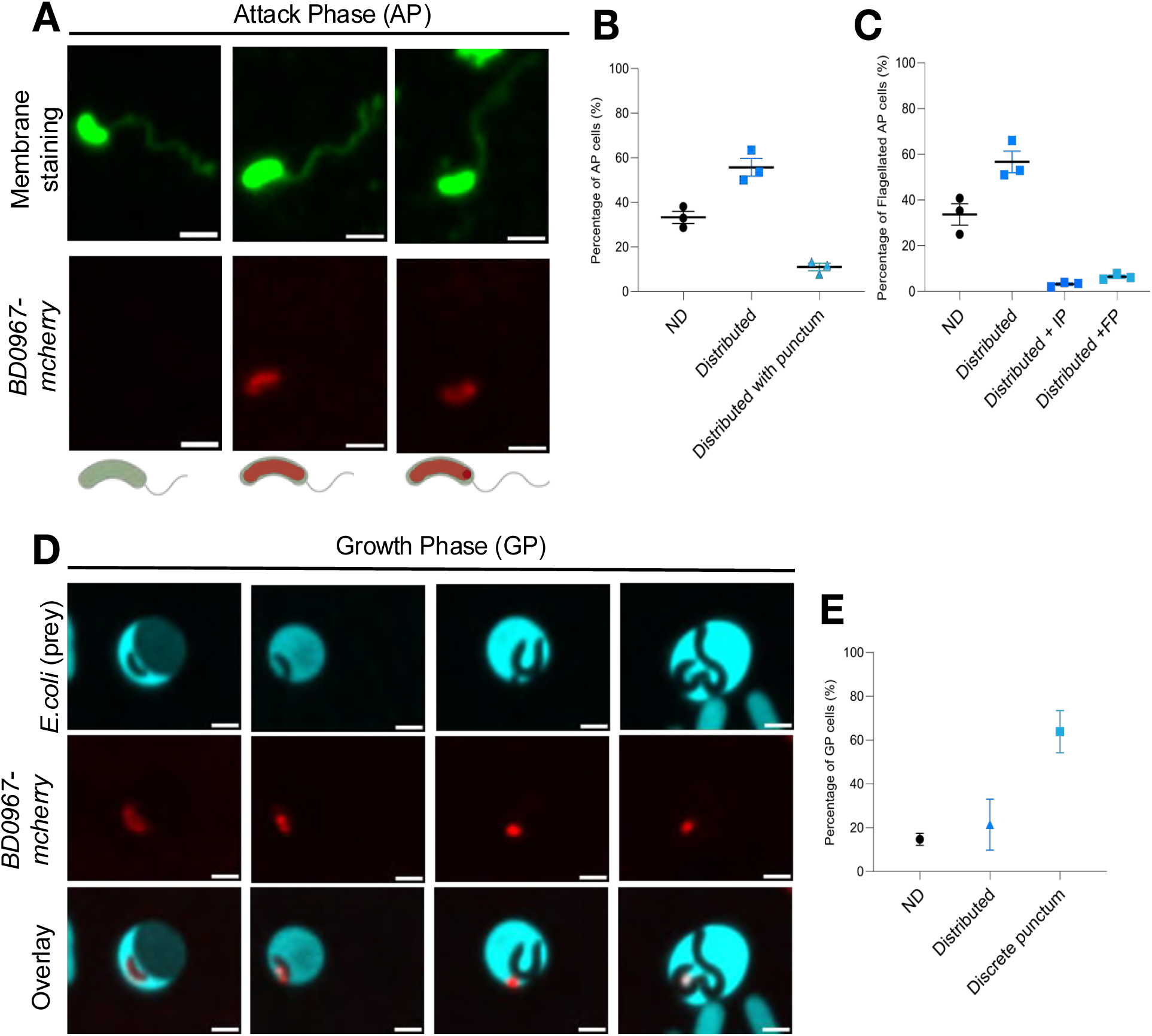
*Bd0967-mCherry* fusion expressed in WT *B. bacteriovorus* HD100 exhibits punctate localisation in bdelloplasts. **(A)** Representative micrographs of attack phase *B. bacteriovorus* HD100 harbouring a C-terminal mCherry (false-coloured red) fusion to Bd0967 at the native locus and stained with Vibrant Dio membrane stain (false-coloured green). Scale bar = 1μm. **(B)** Quantification of cells from (A) plus cells with no flagella detected, n = 296 AP cells from three independent experiments. **(C)** Quantification of flagellated AP cells, n = 156 AP cells from three independent experiments; ND, not detected, IP, invasive pole, FP, flagellar pole. **(D)** Representative micrographs of growth phase cultures two-hours post mixing of prey with predator. *B. bacteriovorus HD100* harbouring the *Bd0967-mCherry* fusion (false-coloured red) can be found within bdelloplasts of *E. coli* DH5α containing plasmid pFED343-Ptac-mTurquoise2 (false coloured cyan). Scale bar = 1μm. **(E)** Quantification of cells from (D), n = 92 GP cells from three independent experiments.

Next, a *B. bacteriovorus* HD100 mutant lacking Bd0967 (Δ*bd0967*) was generated to uncover the role of Bd0967 protease in the predatory life cycle of this bacterium. Although AP cells of the Δ*bd0967* mutant strain had unipolar sheathed flagellum similar to HD100 (SI Figure 12-13), it exhibited several developmental defects. Back-lit microscopy and electron microscopy revealed that the Δ*bd0967* mutant displayed aberrant intracellular growth, inside prey, relative to the wild-type HD100, with irregular morphology and impaired developmental progression (Figure 7 A-C, SI Figure 14). As a next step, by backlighting the *E. coli* prey carrying *pAKF220* plasmid expressing *mNeonGreen* before mixing with *B. bacteriovorus* HD100 and staining nucleic acids with Hoescht (DAPI) stain, we could observe that 15-20% of *E.coli* bdelloplasts, (blue in Figure 7A and green in Figure 7B), formed by the *bd0967* deletion mutant cells rounded up as early as 30 minutes upon prey entry, compared to undetectable levels for bdelloplasts formed by the wild type (Figure 7C). Furthermore, the *bd0967* mutant continued to grow as a larger spherical or aberrantly-shaped cells (black in Figure 7A and blue in Figure 7B) in marked difference to the filamentous growth of the wild-type HD100 predator (Figure 7A-C and SI Figure 15). Consistent with these observations, predation efficiency of the Δ*bd0967* mutant was severely impaired compared to wild-type HD100 as determined by monitoring killing of *E.coli* prey by reduction of OD_600_ over time (Figure 7D-E). Taken together, these findings on timing of prey-invasion and aberrant predator morphology in the *bd0967* mutant suggest that Bd0967 proteolysis of flagellins during flagellar resorption, is involved in the successful progression through the predatory lifecycle.

**Figure 7.**
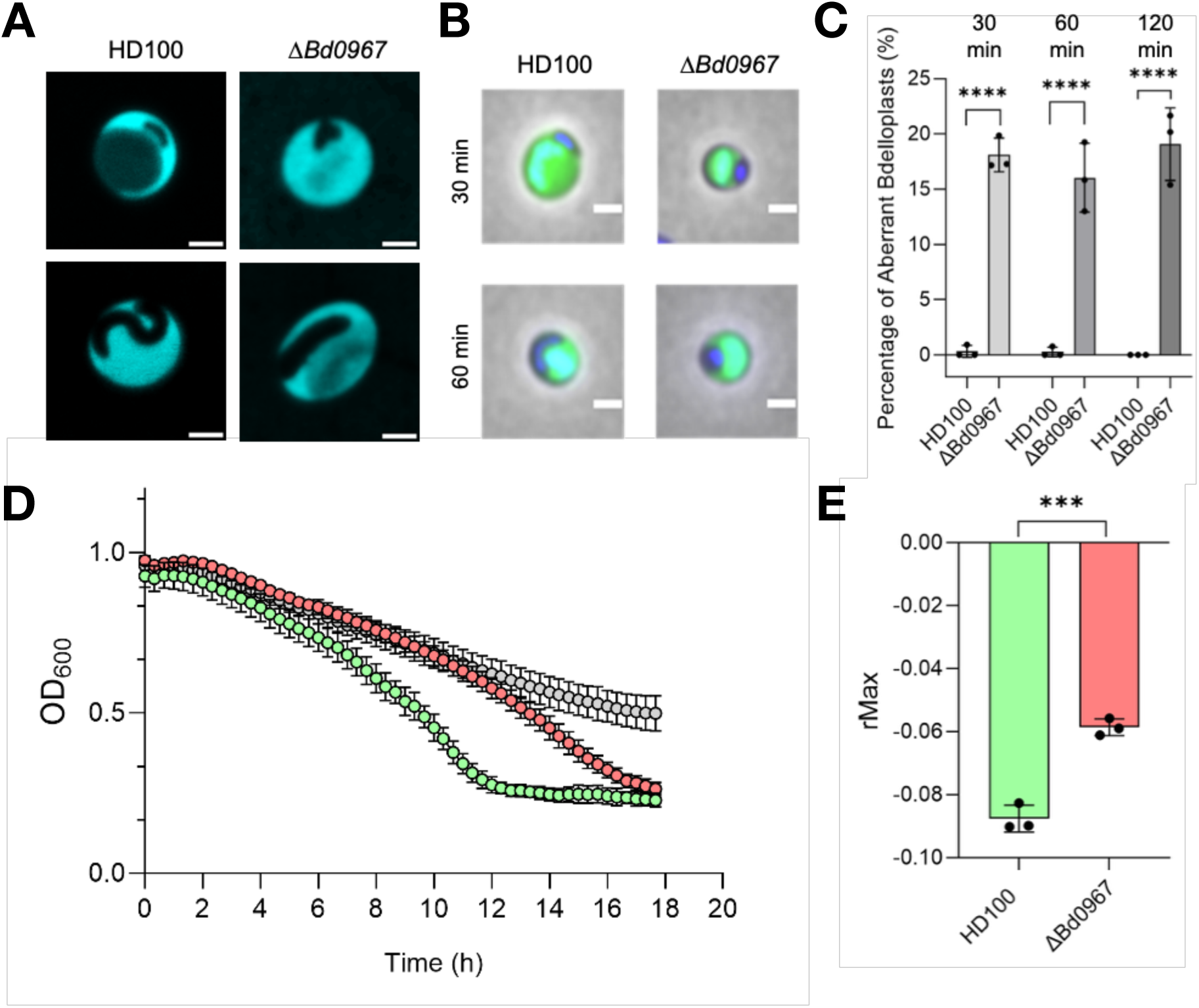
Studying a bd0967 deletion strain shows that Bd0967 contributes to developmental progression. **(A)** Representative micrographs of GP cells (black) in bdelloplasts of E. coli prey (false coloured cyan) visualised by back-lit microscopy. Scale bar = 1μm. **(B)** Epiflourescent and phase contrast merges of wild-type bdelloplasts (HD100) and aberrant growth of Δbd0967 within bdelloplasts at 30 and 60 minutes post-mixing of predator and prey. Prey periplasms are labelled with pMal::mNeon (green) and DNA is stained with Hoechst (blue). Scale bars are 1 μm. **(C)** Quantification of aberrant bdelloplasts at different timepoints post-mixing of predator and prey. Means and independent values are presented from three independent experiments. **** p< 0.0001 by Brown-Forsythe ANOVA. **(D)** Decrease in OD600 of prey E. coli due to predation by Δbd0967 (red) compared to HD100 (green) in microtitre plate assay. **(E)** Measurement of maximum rate of predation (rMax) in microtitre plate assay, measured using curveR. Means and individual values are presented from three independent repeats. *** p=0.0006.

## Discussion

### Protein structure comparisons of Bd0967 and other CTPs

The structure of Bd0967 reveals a divergent CTP architecture in which the conserved S41 protease and PDZ domains are embedded within an extended scaffold, forming a self-enclosed cavity accessed through a proteolytic tunnel. Overall, Bd0967 most closely resembles the structure of Prc from *E. coli*, yet exhibits several distinct structural features that distinguish it from Prc and the more minimal architectures, exemplified by D1P^22^. While the catalytic S41 protease core and PDZ domains are conserved across CTPs, the N-terminal (NTD) and C-terminal (CTD) domains show substantial structural variability. These regions contribute to the overall architecture of each enzyme and appear to be key determinants of functional diversity. For instance, in *Bacillus subtilis* CtpB and *Pseudomonas aeruginosa* CtpA, these domains mediate oligomerisation^2,11^, whereas in *E. coli* Prc and *Chlamydia trachomatis* CT441, they form structural scaffolds that shape substrate access^8,10^. The remarkable architectural diversity within the CTP family, particularly in the N- and C-terminal domains, highlights how these enzymes have evolved to support distinct mechanistic roles while preserving a conserved catalytic core.

The enclosed cavity of Bd0967 resembles a minimal version of the proteolytic compartments found in self-compartmentalising proteases such as the proteasome^27^, ClpP^28^, Lon^29^ and HtrA family members^30^, which form an internal cavity as a result of oligomerisation. This cavity (∼19,500 Å³) is sufficiently large enough to accommodate extended polypeptide segments, such as the C-terminal regions of substrate proteins. This is consistent with the AlphaFold3 models, which predict ∼30-35 residues of the flagellin C-terminus fitting the cavity. The cavity may provide a protected environment that limits solvent exposure and reduces the likelihood of refolding of partially unfolded C-terminal regions. This space may help position and stabilise the substrate polypeptide in a conformation favourable for proteolysis.

The PDZ domain adopts an “open” outward rotated conformation, similar to those observed in other CTPs^2,9,10,12^. Elevated B-factors and conformational sampling observed crystallographically, suggests intrinsic flexibility. This raises the possibility that substrate binding likely induces large scale structural reorganisation of the PDZ domain, similarly to those observed in other CTPs^2^, although such transitions were not directly observed in our structures.

### Mechanism of substrate recognition and turnover

Our findings show Bd0967 interacts with and degrades several *Bdellovibrio* flagellins, with differing efficiencies, where Bd0606 represents the most efficiently processed substrate. We established that the C-terminal interaction with PDZ domain was central to substrate recognition and efficient turnover. Sequence analysis revealed a conserved C-terminal motif amongst *B. bacteriovorus* flagellins (Figure 5B & SI Figure 9). A major exception is Bd0408 (FliC3), which has a slightly longer sequence, primarily due to the addition of 5 residues to the C-terminus. Consistent with the divergence, Bd0408 was the least efficiently degrade substrate from the panel of flagellins. Interestingly, Bd0408 is located at the proximal end and is uniquely essential for functional flagellum assembly^15^.

Structural analysis indicates the flagellin C-terminal motif is well accommodated within the predominantly hydrophobic PDZ peptide-binding pocket. In particular, a basic residue at the P(-3) position (Arg or Lys – consistent motif) models favourably to form a potential electrostatic interaction with D623 of the Bd0967 CTD. This arrangement is strongly supported by AlphaFold3 predictions, which consistently places the C-terminus of the flagellins into the PDZ substrate binding site with high confidence (Figure 5F). Together, the agreement between the consensus motif, structural features of the binding pocket, and computational models supports a mechanism in which Bd0967 recognises substrates through direct interaction with their C-terminal sequences via the PDZ domain, which consequently positions the substrate for cleavage.

Bd0967 degrades Bd0606 through a progressive mechanism characterised by transient accumulation of intermediate fragments. Analysis of degradation patterns by SDS-PAGE revealed a series of reproducible cleavage events, generating products approximately 2–4 kDa smaller than the parent protein before reaching sizes below the resolution limit of the assay (< 10 kDa). This is consistent with Bd0967 acting as an endopeptidase, typical for CTPs. The resulting peptide fragments may then be further processed by other cellular peptidases, such as aminopeptidases and recycled. Consistent with a central role for C-terminal recognition, truncation of the flagellins C-terminus substantially reduces turnover efficiency, while peptide mimics competitively inhibit proteolysis and mutation of a key PDZ residue impairs activity. Nevertheless, truncated substrates are still degraded, indicating that C-terminal recognition enhances substrate capture and positioning within the protease rather than being an absolute requirement for cleavage.

Based on our structural and biochemical data, together with previously described mechanisms for related CTPs^2,10^, we suggest that binding of the exposed flagellin C-terminus to the PDZ domain promotes a conformational change that stabilises an active state of Bd0967 and positions the substrate for cleavage within the protease tunnel. Subsequent cleavage events generate new C-termini that retain features compatible with PDZ-domain recognition, particularly small hydrophobic residues. We therefore propose that flagellin degradation proceeds through iterative cycles of C-terminal recognition and proteolysis, progressively reducing the substrate to smaller fragments. Although our data do not distinguish between a processive mechanism and repeated cycles of substrate dissociation and rebinding, the sequential appearance of discrete intermediates strongly supports a model of progressive substrate processing. This model resembles HtrA-family proteases, where peptide-binding sites cooperate to maintain substrates in a processing-competent state during iterative cleavage^30^.

A key question is how Bd0967 gains access to its substrate *in vivo*. In assembled flagellar filaments, the N- and C-terminal regions of flagellins form a tightly packed helical core buried around the hollow channel within the filament interior^31,32^, making them inaccessible to the folded protease. The internal filament channel (∼20 Å diameter^31,32^), is substantially smaller than the dimensions of Bd0967 (80 x 40 x 50 Å), arguing against direct access to intact filaments. However, recent cryo-electron tomography (cryo-ET) studies, suggest that the *B. bacteriovorus* flagellum is resorbed into the periplasm and subsequently fragmented^20^. Following motor and hook disassembly and filament breakage, the normally buried flagellin terminal regions would become exposed and accessible to Bd0967. Monomeric flagellins in solution (like those that we tested here) are known to have partially unfolded and conformationally dynamic N and C-terminal regions^33^. Following initial filament disruption by physical damage or the action of another different enzyme, exposure of these normally buried termini would therefore generate flexible regions, allowing Bd0967 to engage, recognise and degrade the substrate flagellins via their C-termini. A schematic illustrating the proposed sequence of flagellar resorption, filament destabilisation, and Bd0967-mediated flagellin degradation within the context of the predatory lifecycle is shown in SI Figure 16.

Recognition of flagellins via their terminal regions is well established in other biological contexts. For example, the flagellar chaperones FliS recognises and separates the flagellins N and C-terminal helices during filament assembly^34^. Additionally, exposed C-terminal regions of shed *Salmonella typhimurium* FliC proteins are recognised by host inflammasome components^35^. Together, these observations support the idea that once exposed after flagellum breakage, within the periplasm, the flagellins conformationally dynamic recognition elements would become accessible to Bd0967 PDZ-mediated interaction mechanism.

### Role of Bd0967 and its importance to the lifecycle of *Bdellovibrio*

One of the longstanding mysteries in *B. bacteriovorus* physiology has been the fate of the polar flagellum during transition from the motile attack phase to intracellular growth. *B. bacteriovorus* is a highly motile predator, capable of reaching speeds of 160 μm/s ^15^, driven by its single polar flagellum. This flagellum is essential for prey location during the attack phase but is dispensed of once the predator invades its prey. Early studies suggested the flagellum was shed following predation^19^, whereas later observations indicated it could be retained in a proportion of cells^15^. More recent high-resolution cryo-ET now support a refined model, in which the flagellum is instead resorbed into the periplasm^20^, which represents a rare physiological event, likely reflecting an adaptation to the intraperiplasmic predatory lifestyle of *B. bacteriovorus*. Resorption would permit the recovery of the substantial flagellar biomass, estimated to comprise approximately 10-20,000 flagellin subunits^31^. This could provide a readily accessible source of amino acids during the early stages of prey invasion, before predatory enzymes are ready to consume prey proteins.

Although flagellar assembly has been extensively studied, mechanisms of flagellar disassembly remain comparatively poorly understood across bacteria. In several organisms flagellar loss is coupled to developmental or environmental transitions. For example, *Caulobacter crescentus* uses the ClpAP protease to degrade the flagellar MS ring protein FliF during swarmer-to-stalked cell differentiation, triggering flagellar ejection and the transition to a sessile lifestyle^36,37^. While several γ-proteobacteria eject their polar flagella under nutrients limitation and stress conditions, to conserve energy and halt motility^38^. In contrast, *B. bacteriovorus* appears to internalise its flagellum prior to degradation, representing a highly unusual mode of flagellar remodelling.

Our data provide a direct molecular link between a CTP-family protease and flagellar resorption in *B. bacteriovorus*. Bd0967 carries an N-terminal signal sequence consistent with periplasmic localisation, placing the enzyme in the appropriate compartment to process the resorbed flagella filament. Its ability to selectively degrade multiple *B. bacteriovorus* flagellins *in vitro* identifies a physiologically plausible substrate class for the protease. Furthermore, the highly efficient turnover of flagellins in vitro, is consistent with a biological role in rapidly degrading the large quantity of flagellins upon filament resorption. The localization patterns of the C-terminal mCherry fusion provide spatial context for the enzyme’s function across the predatory life cycle. During the attack phase, the Bd0967-mCherry fusion protein appears diffusely distributed, consistent with a periplasmic or envelope-associated localization in free-swimming predators. In contrast, during growth within the bdelloplast, the fusion protein predominantly forms discrete puncta that are spatially separated from the growing *B. bacteriovorus* filament but remain within the bdelloplast compartment, suggesting association with material in the bdelloplast lumen rather than predator cytoplasmic structures. One interpretation is that these puncta correspond to sites where flagellar remnants (fragments of the filament internalised to the predator’s periplasm through the sheath retraction mechanism) accumulate following resorption where they serve as concentrated substrates sites. Consistent with a functional role for Bd0967 at these sites, deletion of the protease gene results in aberrant bdelloplast morphology and slower predation dynamics. These phenotypes suggest failure to efficiently degrade resorbed flagellin, disrupts the confined bdelloplast environment or the polarity organisation of the developing and elongating Bdellovibrio cell filament which later septates into progeny along its length, and delays the physiological transition from motility to intracellular growth.

In *B. bacteriovorus*, flagellar resorption and subsequent degradation by Bd0967 may confer multiple advantages. First, it would conserve energy and recycle valuable amino acid resources by degrading resorbed flagellin subunits rather than discarding them, providing an immediate source of amino acids from the predator’s own flagellar proteins prior to the extensive digestion of prey macromolecules. Second, it would prevent the release of free flagellin into the extracellular environment. Flagellin is a potent pathogen-associated molecular pattern (PAMP) recognised by Toll-like receptor 5 (TLR5), triggering strong innate immune responses including NF-κB activation and pro-inflammatory cytokine production^39,40^. For a predatory bacterium which may reside within mammalian hosts, or may be similarly sensed by Protozoa in natural soil and water environments, minimising the immunogenic footprint of its lifecycle transitions could be biologically advantageous. Third, flagellar resorption and degradation may function as a committed signalling event that reinforces the transition from the motile attack phase to the non-motile, replicative growth phase by remodelling the old flagellar pole of the invader.

CTPs serve as important regulators of developmental transitions in diverse bacteria^1^ and the link between generalised flagellar dynamics and cell cycle control is well established across multiple organisms, including CtrA-regulated flagellar biogenesis in *C. crescentus*^41^. Our work provides a direct molecular link between a CTPs and flagellar remodelling in *B. bacteriovorus*, demonstrating that Bd0967-mediated flagellin degradation is coupled to normal lifecycle progression. Taken together, the biochemical specificity of Bd0967 for flagellin, its localisation during intracellular growth, and the developmental defects arising from its loss, support a model in which Bd0967 functions as a dedicated component of the flagellar resorption programme. More broadly, these findings expand the known functional repertoire of CTPs beyond their canonical roles in protein quality control and peptidoglycan remodelling to include selective flagellar remodelling during bacterial developmental transitions.

## METHODS

### Materials, chemicals and general

Chemicals used in this investigation were purchased from Fisher and Merck. Peptides were kindly provided by the Peacock group (University of Birmingham) and Singh group (University of Liverpool). The synthesis of these peptides is found in the supplementary material, methods section. Primers used for cloning, were purchased from IDT or Sigma-Aldrich. A list of primers used in this study is provided in Supplementary Table 1 and a list of plasmids is provided in Supplementary Table 2. Kinetic data were fitted using GraphPad Prism. SDS-Page gels were analysed by ImageJ.

### Strains, media, and growth conditions

Strains used in this study are listed in Supplementary Table 3. *E. coli* S17-1 was routinely grown on YT agar plates and in YT broth for 16 hours at 37°C with shaking at 200 rpm. *B. bacteriovorus* were grown as a predatory culture on stationary phase *E. coli* S17-1 prey for 24 hours at 29°C in Ca/HEPES buffer (5.94g/l HEPES free acid, 0.284 g/l calcium chloride dihydrate, pH 7.6), or on YPSC overlay plates as plaques within a lawn of *Escherichia coli* S17-1 prey, as described previously^42^.

### Generation of markerless deletion mutants

To construct a markerless gene deletion of *bd0967*, 1000bp of DNA upstream and downstream of the gene of interest were cloned into the suicide vector pk18*mobsacB* by Gibson assembly (Gibson et al., 2009) using the NEBuilder HiFi DNA assembly cloning kit (New England BioLabs). Primers used are listed in Supplementary Table 1. To return the *bd0967* gene to the mutant creating a complement strain, the external primers 967KOupF and 967KOdownR were used to clone the gene with flanking DNA. Gene deletion or complementation vectors were introduced into *B. bacteriovorus* by conjugation (using donor *E. coli* S17-1 strains) and subsequently cured of the donor plasmid by sucrose suicide counter-selection, resulting in the integration of the constructs via double crossover homologous recombination. This process was described previously^43^. All gene deletions were verified by Sanger sequencing.

### Generation of Bd0967-mCherry fusion

For construction of the fluorescently tagged bd0967, mCherry was fused to the C-terminus of bd0967. The stop codon of bd0967 was removed, and a linker (amino acid sequence: DILEL) was inserted between the gene and the mCherry sequence. This linker was used in a previous study in *V. parahaemolyticus* (EE Arroyo-Pérez, et.al., 2021). For conjugation of the plasmid carrying the bd0967–mCherry fusion into B. bacteriovorus, *E. coli* carrying the desired plasmid was grown overnight in 10 ml PY liquid culture, pelleted, and resuspended in 100 µl PY. *B. bacteriovorus* cultures were filtered (50 ml lysates after ∼48 h incubation), centrifuged at maximum speed for 20 min, and resuspended in 100 µl HEPES/CaCl₂. A sterile filter paper was placed in the center of a PY plate, onto which 100 µl of E. coli and 100 µl of filtered *Bdellovibrio* were added. Plates were incubated overnight at 30 °C. The following day, the filter paper was washed in 1 ml HEPES/CaCl₂. The collected cells were mixed with 200 µl of E. coli S17-1 (kanamycin-resistant) and 5 ml of 0.6% YPSC soft agar containing kanamycin (final concentration 50 µg/ml). The mixture was poured onto 1% YPSC bottom agar plates also containing kanamycin (50 µg/ml) and incubated at 30 °C for 3–5 days until plaques appeared. Plaques were picked for further propagation and screened for mCherry insertion. The resulting strain is merodiploid and kanamycin resistant.

### Microscopy of the BD0967-mcherry during attack phase, bdelloplast development (with backlit fluorescent microscopy) and quantification

For the membrane staining, we used the dye Vybrant™ DiO Cell-Labeling Solution (ThermoFisher, Cat. No. V22886). Lysates of B. bacteriovorus (10ml) were grown overnight at 30 °C with constant shaking. Lysates were first filtered and centrifuged at 9,000 rpm for 20 min to pellet the cells. The supernatant was discarded, and the pellet was resuspended in fresh HEPES/CaCl₂ to the original volume (10 ml). The resuspended lysate was allowed to rest at room temperature for 1–2 h. To minimize flagellar breakage, pipette tips were trimmed prior to handling. Aliquots of 1 ml lysate were then centrifuged at 15,000 rpm for 20 min, the supernatant was removed, and the pellet was incubated for 20 min at 37 °C in HEPES/CaCl₂ containing the membrane dye DiO (5 µl dye per ml of medium), using pre-warmed HEPES/CaCl₂ as recommended by the manufacturer. Cells were subsequently centrifuged at 15,000 rpm for 5 min, washed once with warm HEPES/CaCl₂, and centrifuged again for 5 min. The final pellet was resuspended in 20 µl fresh HEPES/CaCl₂, and 10 µl was used for microscopy. Coverslips were pre-coated with poly-L-lysine prior to imaging.

Backlit fluorescence microscopy of *B. bacteriovorus* HD100 expressing bd0967–mCherry was performed as previously described^44^. Fluorescent E. coli prey expressing mTurquoise2 were induced with IPTG (0.1 mM). Overnight E. coli cultures (we used just 500 µl) were mixed in HEPES/CaCl₂ with 1 ml of B. bacteriovorus (from a filtered 10 ml lysate resuspended in 2 ml). Lysates were incubated at 30°C with constant shaking. Samples were collected at time points T0–30 min and T2–5 h. At each time point, 1 ml of lysate was centrifuged and resuspended in 30 µl of fresh HEPES/CaCl₂, and 10 µl was used for microscopy. Coverslips were pre-coated with poly-L-lysine prior to imaging. All microscopy images were taken using the inverted confocal microscope Leica Stellaris 5. Images were then processed using Fiji. Quantification of mcherry puncta was done manually, and values were graphed using GraphPad.

### Phase-contrast and epifluorescence microscopy

Approximately synchronous predation of *E. coli* S17-1 by *B. bacteriovorus* strains was prepared by combining a 10x concentrated *B. bacteriovorus* predatory culture with *E. coli* S17-1 standardised to an OD_600_ of 1.0 and Ca/HEPES at a ratio of 5:4:3 respectively as described previously^45^. Development of the predator within the bdelloplast was monitored at 30-, 60- and 120-minutes post mixing of predator and prey by withdrawing 19 μl of the culture adding to 1 µl of 10 µM Hoescht 33342 (DAPI) and immobilising 10 µl on a thin 1% Ca/HEPES buffer agarose pad. Cells were visualised with a Nikon Ti-E inverted epifluorescence microscope equipped with a Plan Apo 100x Ph3 oil objective lens (NA: 1.45) and the following filters GFP filter (for mNeonGreen; excitation: 460-500 nm, emission: 515-530nm): DAPI (excitation: 340-360 nm, emission: 435-485 nm). Images were acquired on an Andor Neo sCMOS camera with Nikon NIS software and analyzed using ImageJ (FIJI 1.52n).

### Image analysis

Images were manipulated with ImageJ (FIJI distribution 1.52n) software using the sharpen and smooth tools, and by duplication of region of interest (ROI) for presentation. Images were analysed using the MicrobeJ plugin (version 5.13j) for the ImageJ^46^ which automates detection of bacteria within an image. Attack phase *B. bacteriovorus* cells were detected by the medial axis method with the following parameters: area-0.3-2 µm^2^, length 0.5-4 µm, width 0.2-0.85 µm, circularity 0- 0.95 µm and all other parameters as default. Any aberrantly shaped cells not detected by these parameters were manually counted.

### Microscopic Statistical analysis

Statistical analysis was performed in Prism 10.6.1 (GraphPad). Data were first tested for normality with the Shapiro-Wilk test and then analysed using the appropriate statistical test accordingly. The number of biological repeats and the statistical test applied are described within each figure legend.

### *B. bacteriovorus* predation on *E. coli* in liquid culture

Assays were based on, and modified from, those detailed in Remy, O., et al.^47^. In summary, *B. bacteriovorus* strains were grown predatorily (as above). PFU (Plaque Forming Unit) inputs for each strain were matched using SYBR Green DNA stain, to ensure equal titres of *B. bacteriovorus* for each strain were used as starting inputs. Briefly, *B. bacteriovorus* were incubated with SYBR Green dye for 90 minutes (300 rpm double orbital, in darkness, in triplicate), before fluorescence was measured using a FLUOStar Omega Plate Reader (BMG Labtech) (Excitation: 485 nm, Emission: 520 nm, Gain: 800), with fluorescence values being interpolated into relative PFU/ml counts using a PFU:SYBR Green Fluorescence correlation curve. *E. coli* S17-1 cells were grown as above, and subsequently backdiluted to OD_600_ 1.0 (approximately 1 × 10^9^ CFU/ml) in DNB (Dilute Nutrient Broth). Predatory cultures (containing approximately 1 × 10^9^ CFU/ml *E. coli* prey and 1 × 10^8^ PFU/ml *B. bacteriovorus bacteriovorus* WT or mutant) were inoculated in triplicate into a black OptiPlate (Corning), along with media only, *B. bacteriovorus* only (no prey) and prey only (no *B. bacteriovorus*) controls. The optical density at 600nm (OD_600_) of the predatory culture was measured every 20 minutes, for 18 hours (200 rpm, double orbital, in triplicate) to give a prey survival curve, where a drop in OD_600_ is indicative of successful predation and prey lysis, because prey cells but not predators, are large enough to produce an optical density at 600nm. OD_600_ data was exported to Excel 2016 and then analysed using CurveR, according to the method documented in Remy et al. to analyse prey cell lysis and predation dynamics. R_max_ indicates the maximum rate of prey cell lysis.

### Cloning, expression and purification of flagellin and Bd0967 proteins

For this investigation a soluble construct of Bd0967 comprising residues 25-673 was used, lacking the predicted N-terminal signal peptide region (residues 1-24). This gene construct was cloned into pET-28a, following an N-terminal thrombin cleavable His-6 tag, using a restriction free cloning method. Bd0606 was cloned as a full-length construct (residues 1-277) into pET28a, as for Bd0967. Mutants (R299M & S457A) and truncations (LIG & RLIG) of these constructs were generated using a standard site directed mutagenesis protocol. Constructs and mutations were confirmed by sangar sequencing before transformation into the *E. coli* expression strains.

Bd0967 constructs were transformed into SHuffle® T7 by NEB, to assist disulphide bond formation, whilst Bd0606 constructs were transformed into Bl21(DE3) cells. Transformed cells were grown at 37 °C (shaken at 180 rpm) in LB media (10 g/L NaCl, 10 g/L tryptone, and 5 g/L yeast extract) supplemented with 100 μg/ml Kanamycin until an OD_600_ of 0.8 was reached. Gene expression was induced with 1 mM IPTG and incubated over night at 16 °C (shaken at 180 rpm). Cells were harvested by centrifugation at 8855 g (Beckman JLA 8.1000 rotor) for 10 min and the pellets stored at −20 °C.

Cells were re-suspended in fresh lysis buffer (50 mM HEPES pH 8.0, 20 mM Imidazole pH 8.0, 500 mM NaCl) and incubated with ∼1 mg/ml lysozyme for 30 min at 4 °C. Cells were lysed using sonication and insoluble cell debris was removed by centrifugation for 30 minutes at 33,000 g (Beckman JA 25.50). The supernatant was loaded onto a 5 ml HisTrap HP column (GE Healthcare) pre-equilibrated with lysis buffer. The column was washed with 20 column volumes (CV) of lysis buffer and 10 CV 10% elution buffer solution (50 mM HEPES pH 8.0, 500 mM NaCl, 300 mM Imidazole pH 8.0) to remove non-specific interacting proteins. 10 CV of elution buffer was passed over the column to elute the His_6_-tagged proteins. Fractions containing pure protein of interest (e.g. Bd0967 / Bd0606), evaluated by SDS-PAGE, were pooled and dialysed over night against 2 L of dialysis buffer (20 mM HEPES pH 7.5, 300 mM NaCl, 5 mM MgCl_2_). Pooled & dialysed samples were concentrated to ∼5 ml and injected onto a 26/600 S200 size exclusion chromatography column for polishing. Fractions were analysed by SDS-PAGE to ensure purity and samples containing proteins of interest were pooled and concentrated to ∼10 mg/ml for further experiments. Protein concentration was deduced from A280 readings using a thermoscientific NanoDrop One instrument for Bd0967.

Bradford assays using thermoscientific BioMate 160 UV-visible spectrophotometer was used to determine protein concentration of Bd0606, which lacks tryptophan residues.

### Crystallisation, data collection and structure determination

Crystals were first grown at 18°C using sitting drop vapour diffusion technique, with a drop size of 0.3 μl in a ratio of 1:1, reservoir:protein solution, using ttplabtech mosquito HTS nanolitre liquid handler. Initial crystals grown in Morpheus G4 were small fibrous needles, which were used as a seed stock in successive rounds of random matrix microseeding^48^ (rMMS). rMMS trials set up using a 3:2:1 ratio of protein:reservoir:seed stock, with the initial seed stock diluted 1 in 10 prior to use. Seeding resulted in improved crystal growth and diffraction quality across several conditions. Larger rod-shaped crystals of WT Bd0967 obtained in the Morpheus G4 condition, following seeding, were used to collect the high-resolution data. Bd0967_S457A_ crystals were grown in Morpheus G12, which had been seeded with WT crystals. A full list of crystallisation conditions can be found in Supplementary table 4. Crystals in mother liquor were flash cooled in liquid nitrogen and stored under liquid nitrogen until data collection.

Diffraction data were collected at the Diamond Light Source (Oxford, UK) on beamlines I04 & I04-I. Data reduction and processing were carried out using automated pipelines with XDS and the xia2 suite. The structure of WT Bd0967 was solved by molecular replacement, using a chainsaw-edited model derived from chain A of PDB: 4QL6 (CT441)^8^. Chainsaw was used to prune the sidechains of residues to match those of Bd0967 in a sequence alignment^49^. The initial model was improved through Phenix AutoBuild and manual model building in Coot. The Bd0967_S457A_ structure was solved using the structure of WT bd0967 as the search model in PHASER. Subsequent model building and refinement were performed iteratively in Coot and Phenix. Metal ion selection and placement was guided by electron density, coordination and was limited to elements found in crystallisation conditions. Co-crystallised peptides were modelled and refined as poly-alanine chains; although electron density indicated the presence of larger side chains, these could not be assigned unambiguously at the achieved resolution. Crystallographic data are shown in Supplementary Table 5.

### Cavity calculation

The internal cavity of Bd0967 was assessed using the ‘Find Cavities’ tool in ChimeraX 1.10.1, developed by KVFinder^21^. The cavity was calculated with the following parameters: grid spacing = 0.6 Å; inner probe radius = 2.5 Å; outer probe radius = 10 Å; Exterior trim distance = 5 Å; to appropriately represent the internal cavity of Bd0967. The complete set of calculated values for the cavity are as follows: volume = 13952 Å^3^, surface area = 3979 Å^2^, average depth = 7.49 Å, Maximum depth = 16.26 Å.

### AlphaFold predictions and modelling

AlphaFold3 was used to generate protein–protein interaction models between Bd0967 and *Bdellovibrio bacteriovorus* flagellins (Bd0408, Bd0410, Bd0604, Bd0606, Bd3052, and Bd3342). The amino acid sequences of Bd0967 (secreted form) and each flagellin were used as input. For each Bd0967–flagellin interaction, multiple models were generated and ranked according to the model confidence metrics provided by AlphaFold3. AlphaFold3 was also used to model interaction between Bd0967 and the panel of recombinantly expressed *Bdellovibrio bacteriovorus* proteins (Bd0108, Bd0314, Bd1833, Bd1996, Bd2924) and to model the interaction between Bd0606 and other *Bdellovibrio bacteriovorus* C-terminal processing proteases (Bd0169, Bd1239 and Bd3534).

Predicted structures were analysed using the per-residue confidence score (pLDDT) and predicted aligned error (PAE) to assess model reliability, with particular focus on the interface between Bd0967 and the flagellin C-terminal region. Structural models were visualised and analysed using Coot and ChimeraX. The interface regions and putative binding modes were interpreted qualitatively, with emphasis placed on consistently observed features across independent predictions, such as the PDZ and C-terminus interactions.

### Ni-NTA pulldown assay

A pulldown assay was performed to identify interacting proteins & substrates of Bd0967. *Bdellovibrio bacteriovorus* HD100 cells were grown to a cell density of 2.5×10^8^ cells/ml and then harvested by centrifugation 6000 RPM for 20 minutes at 4°C in a JLA-10.5 rotor. The supernatant was discarded and the cell pellet was resuspended in ice cold lysis buffer (20mM Na Phosphate pH 7.4, 40 mM Imidazole pH 8.0, 300 mM NaCl) supplemented with a cOmplete protease inhibitor tablet (Roche) and lysozyme at 1 mg/ml. Resuspended cells were fractioned and lysed by sonication in cycles of 20 s on/off, for 5 minutes. Insoluble material was then removed by centrifugation at 38000 g for 45 minutes at 4 °C. The resulting soluble fraction was carefully removed and stored at 4 °C for pulldown assays. The remaining pellet was resuspended in ice cold lysis buffer supplemented with 2% Triton X-100, followed by centrifugation at 38000g for 10 minutes to remove cell debris and prepare the membrane fraction. Both soluble and membrane fractions were prepared and used in same-day pulldown assays.

The pulldown assay was prepared by: gently pelleting a 200 μl slurry of Qiagen Ni-NTA beads (2500 g for 2 minutes at 4°C); equilibrating beads with lysis buffer by preforming three wash steps (addition of 500 μl lysis buffer); final resuspension of beads into 50 μl lysis buffer. The pulldown assay was performed using the inactive Bd0967_S457A_ mutant, to prevent unwanted protein degradation, which may negatively affected detection of interacting proteins. Bd0967_S457A_ was diluted with wash buffer (20mM sodium phosphate pH7.4, 300mM NaCl, 40mM imidazole) to 150 μg/ml and 1ml was then incubated with 50 μl of prepared bead solution for 30 minutes at 4°C by end over end rotation. A lysis buffer only control (C1 = beads plus 1 ml lysis buffer), was also prepared side by side. After incubation the beads were pelleted and washed three times with wash buffer to remove unbound proteins.

Bd0967-bound beads and control beads (C1) were then incubated for 2 hours with 5ml of soluble or membrane fractions at 4°C, by end over end rotation. Another control, C2, where Bd0967-bound beads were incubated with lysis buffer for 2 hours was run alongside those incubated with protein fractions. After incubation, unbound proteins were removed by performing 3 wash steps with wash buffer. Washed beads were pelleted, resuspended in 100 μl 1x Laemeli buffer and boiled at 100°C for 15 minutes to prepare samples for SDS-PAGE analysis. Gels were stained and imaged to visualise interacting proteins. Gels with visible interacting proteins were dissected into lanes and stored at −20°C, prior to in gel trypsin digest and mass spectrometry analysis.

### LC-MS protein identification (nanoLC-ESI-MS/MS Analysis)

Reversed phase chromatography was used to separate tryptic peptides prior to mass spectrometric analysis. Two columns were utilised, an Acclaim PepMap µ-precolumn cartridge 300 µm i.d. x 5 mm 5 μm 100 Å and an Acclaim PepMap RSLC 75 µm x 25 cm 2 µm 100 Å (Thermo Scientific). The columns were installed on an Ultimate 3000 RSLCnano system (Dionex). Mobile phase buffer A was composed of 0.1% formic acid in water and mobile phase B 0.1 % formic acid in acetonitrile. Samples were loaded onto the µ-precolumn equilibrated in 2% aqueous acetonitrile containing 0.1% trifluoroacetic acid for 8 min at 10 µL min^−1^ after which peptides were eluted onto the analytical column at 300 nL min^−1^ by increasing the mobile phase B concentration from 4% B to 25% over 37 min, then to 35% B over 10 min, and to 90% B over 3 min, followed by a 10 min re-equilibration at 4% B.

Eluting peptides were converted to gas-phase ions by means of electrospray ionization and analysed on a Thermo Orbitrap Fusion (Q-OT-qIT, Thermo Scientific)^50^. Survey scans of peptide precursors from 375 to 1500 *m*/*z* were performed at 120K resolution (at 200 *m*/*z*) with a 2 × 10^5^ ion count target. Tandem MS was performed by isolation at 1.2 Th using the quadrupole, HCD fragmentation with normalized collision energy of 33, and rapid scan MS analysis in the ion trap. The MS^2^ ion count target was set to 5×10^3^ and the max injection time was 200 ms. Precursors with charge state 2–6 were selected and sampled for MS^2^. The dynamic exclusion duration was set to 45 s with a 10 ppm tolerance around the selected precursor and its isotopes. Monoisotopic precursor selection was turned on. The instrument was run in top speed mode with 2 s cycles.

### Data Analysis

The raw data were searched against *Bdellovibrio bacteriovorus and Escherichia coli* databases and the common contaminant database from MaxQuant (https://www.maxquant.org/). MaxQuant software (version 1.5.5.3) was used for protein identification. Peptides were generated from a tryptic digestion with up to two missed cleavages, carbamidomethylation of cysteines as fixed modification and oxidation of methionine as variable modification. Scaffold (TM, version 4.4.5, Proteome Software Inc.) was used to validate MS/MS based peptide and protein identifications. Peptide identifications were accepted if they could be established at greater than 95.0% probability by the Scaffold Local FDR algorithm. Protein identifications were accepted if they could be established at greater than 99.0% probability and contained at least 2 identified peptides. Proteins that contained similar peptides and could not be differentiated based on MS/MS analysis alone were grouped to satisfy the principles of parsimony.

### Degradation assay

Bd0967 proteolytic activity was assessed using a discontinuous, gel-based degradation assay with soluble purified Bd0606 as substrate. Reactions were carried out under standard conditions at 37 °C with agitation in 50 mM HEPES (pH 7.5), 250 mM NaCl, 5 mM MgCl_2._ Assays were initiated by adding Bd0967 (1 μM final) to reaction mixtures containing Bd0606 (100 μM final); total volumes varied depending on the experiment.

Reactions were quenched by addition of 4× Laemmli sample buffer (LSB) followed by incubation at 100 °C for 10 minutes. For analysis, 10 μL of each quenched sample (from 15 μL reaction + 5 μL 4× LSB) was loaded onto SDS–PAGE gels and operated under standard running conditions. Gels were stained and imaged using a Bio-Rad ChemiDoc MP system.

For standard degradation assays, reactions were quenched after 60 min, which corresponded to the reaction endpoint for WT Bd0967 under these conditions. For time-course assays, a master mix of sufficient volume for all time points was prepared, and 15 μL aliquots were withdrawn at the indicated times (1–120 min) and quenched as described above. Due to experimental set up it was not possible to get an accurate 0-min timepoint.

For substrate denaturation assays, potential substrates (lysozyme and BSA) were added to standard reaction buffer at a concentration of 1 mg/ml (final concentration). Heat denaturation at 100 °C for 10 mins was performed in the reaction tube, to prevent unrelated loss of substrate mass to transferring sample, which may have confounded interpretation. Sample were left to equilibrate to room temperature before the addition of Bd0967 to a final concentration of 1 μM, at which point samples were transferred to 37 °C shaking incubator. After 60 minutes the samples were quenched and prepared for SDS-PAGE as above. Heat denatured samples were run alongside native protein and a +ve control (native Bd0606).

Peptide inhibition assays were performed under standard conditions with peptides added to a final concentration range of 0–12 μM. Reactions were incubated for 90 min at 37 °C, quenched, and analysed by SDS–PAGE as described. Inhibitors (Inhibitor-1, NAMPNSALRLIG and Inhibitor-2, NAMPNSALRIG) were kindly synthesised through collaborations with the Peacock and Singh groups. Details of peptide synthesis and purity analysis are provided in the Supplementary Information (SI Figures 17 & 18).

All enzyme assays were performed on at least three independent occasions using the same protein preparations.

### Gel Analysis and Data fitting

SDS-PAGE gel images were analysed in ImageJ^51^ (FIJI software) using the programs gel analysis tool. Lanes were assigned manually with the rectangle selection tool to encompass the full migration range of each sample. Band intensity profiles were generated and the peak areas corresponding to Bd0967 and Bd0606 were integrated. Intensity values (arbitrary units, AU) were exported, then plotted and analysed in GraphPad Prism. Peptide degradation time-course data were fit to the following decay models. WT Bd0967 degradation of WT Bd0606 substrate followed a one phase decay model (Equation 1). Whereas degradation of ΔLIG and ΔRLIG substrates showed a clear initial plateaux and was fit to the plateau followed by one phase decay model (equation 2).

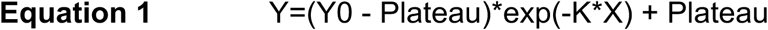

Y is the remaining substrate at a given time (X); Y0 is the Y value when X (time) is zero; Plateau is the Y value at infinite times; K is the observed rate constant.

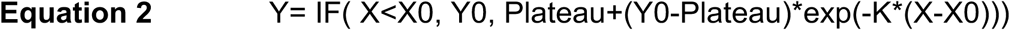

Y is the remaining substrate at a given time (X); X0 is the time at which decay begins; Y0 is the average Y value up to time X); Plateau is the Y value at infinite times; K is the observed rate constant.

Supplementary Table 6 details fitted data for each substrate species.

### Circular Dichroism

Circular dichroism (CD) spectroscopy was used to assess the secondary structure and folding of purified flagellin proteins. Far-UV CD spectra were recorded on a Jasco J-1100 CD spectrometer, controlled using the Jasco spectra manager 2.0 software version 2.0. Measurements were carried out at 20 °C using a 1 mm pathlength quartz cuvette. WT Bd0606 was used to optimise buffer conditions and protein concentration for CD measurements. Protein samples were prepared at a concentration of ∼0.25 mg/ml in 20 mM sodium phosphate, pH 8.0 to limit buffer absorbance affects.

Spectra were collected over a wavelength range of 190–280 nm with a bandwidth of 1 nm, a scan speed of 100 nm/min, and averaged over ten accumulations. Buffer spectra were recorded under identical conditions and subtracted from sample spectra. Raw ellipticity data were converted to mean residue ellipticity (MRE) to allow comparison between proteins of different lengths.

### Sequence alignments and conservation analysis

Multiple sequence alignments of *B. bacteriovorus* flagellins were performed using Clustal Omega with default parameters. Alignments were visualised and manually inspected to assess conservation across the full-length sequences and, in particular, within the C-terminal region. The image of the alignment was generated using ESPript3.

To analyse positional conservation at the C-terminus, aligned sequences were trimmed to the final residues and WebLogo3 ^52^ was used to generate sequence logos, which represent the frequency and conservation of amino acids at each position^53^.

The Genome.jp database was used to search for candidate proteins containing similar C-terminal sequences, using the proteome of *B. bacteriovorus* HD100. A motif-based search was performed using a permissive consensus motif ([LMIV]-[RKHMG]-[LIV]-[ILVMFGA]-[GAN]->) to capture a broader set of potential matches within the proteome.

### Statistical analysis

All data are presented as mean ± standard error of the mean and were obtained from ≥3 independent experiments with total sample numbers provided in the figure legends. Statistical significance was evaluated with GraphPad Prism software, using a one-sample t-test. Significant difference assessed through an unpaired t-test, ****P < 0.0001.

## Data availability

Structural data were deposited in the PDB and are available under accession numbers 30XH & 30XJ. All other data are contained in the main manuscript and supplementary information. Raw data accompanying figures is available for download on figshare (https://doi.org/10.6084/m9.figshare.32546355). All Alphafold3 models predicting CTP and substrate interactions have been shared using figshare - https://doi.org/10.6084/m9.figshare.32567625. All materials and reagents are available from the corresponding authors.

## Acknowledgements

We thank the School of Life Sciences Imaging Department (SLIM) at the University of Nottingham for assistance with electron microscopy. We are grateful to the WPH Proteomics RTP Facility at the University of Warwick, particularly Dr Cleidi Zampronio, for performing and assisting with analysis of mass spectrometry experiments. We thank the Protein Analysis Facility at the University of Liverpool, particularly Dr Igor Barsukov, for access and assistance with circular dichroism experiments and data analysis. We thank Professor Kim Hardie for hosting C.L. and supporting completion of the analysis presented in this study.

P.R., C.L., R.T., S.C., A.L.L., and R.E.S. were supported by a Wellcome Trust Investigator Award in Science (209437/Z/17/Z to R.E.S. and A.L.L.). CH was supported by a BBSRC Midlands Integrative Biosciences Training Partnership (MIBTP) PhD studentship (1500753), and a Royal Society Research grant (RG\R1\251063). SM and A M-L were funded by the University of Chicago Start-up funds to S.M.. K.A.H. was supported by a PhD studentship funded by the EPSRC CDT in Topological Design (EP/Z533452/1).

## Author contributions

**Conceptualisation and study design:** C.J.H., S.M. and A.L.L..

**Investigation:** C.J.H. performed the structural, biophysical, biochemical and substrate-identification studies, including protein production (Bd0967 & flagellins), crystallography, pulldown experiments, enzymatic characterisation, and substrate-recognition analyses. A.M.-L. performed fluorescence microscopy, mutant phenotyping and predation assays. C.L. designed the *bd0967* deletion strategy and carried out microscopy, image analysis, predation assays and phenotypic characterisation of mutant strains. R.T. constructed and verified the Δ*bd0967* mutant. S.C. performed preliminary screening of additional flagellin constructs. P.R. generated the complementation strain and performed microscopy experiments. K.H. and A.P. synthesised the peptides used in the inhibition studies, while O.D. synthesised a peptide used in preliminary experiments.

**Methodology and supervision:** A.F.A.P. and I.S. supervised peptide synthesis. S.M. supervised the fluorescence microscopy and mutant phenotyping studies. A.L.L. supervised the structural and biochemical studies and overall project. R.E.S. supervised microscopy and mutant phenotyping studies performed at the University of Nottingham.

**Data analysis and interpretation:** C.J.H., A.M.-L., C.L., S.M., L.S. and A.L.L.

**Writing:** C.J.H. wrote the original manuscript draft. C.J.H., C.L., S.M., R.E.S. and A.L.L. reviewed and edited the manuscript.

All authors approved the final manuscript.

## Notes

### Competing Interest Statement

The authors have declared no competing interest.

### Summary of Updates

Update the manuscript -> Figures within main text Inclusion of Supplementary Information

## References

1 Sommerfield, A. G. & Darwin, A. J. Bacterial Carboxyl-Terminal Processing Proteases Play Critical Roles in the Cell Envelope and Beyond. Journal of Bacteriology 204, doi:10.1128/jb.00628-21 (2022).

2 Mastny, M. et al. CtpB Assembles a Gated Protease Tunnel Regulating Cell-Cell Signaling during Spore Formation in Bacillus subtilis. Cell 155, 647–658, doi:10.1016/j.cell.2013.09.050 (2013).

3 Singh, S. K., Parveen, S., SaiSree, L. & Reddy, M. Regulated proteolysis of a cross-link-specific peptidoglycan hydrolase contributes to bacterial morphogenesis. Proc Natl Acad Sci U S A 112, 10956–10961, doi:10.1073/pnas.1507760112 (2015).

4 Srivastava, D. et al. A Proteolytic Complex Targets Multiple Cell Wall Hydrolases in Pseudomonas aeruginosa. mBio 9, doi:10.1128/mBio.00972-18 (2018).

5 Carroll, R. K. et al. The lone S41 family C-terminal processing protease in Staphylococcus aureus is localized to the cell wall and contributes to virulence. Microbiology 160, 1737–1748, doi:10.1099/mic.0.079798-0 (2014).

6 Roy, R., You, R.-I., Lin, M.-D. & Lin, N.-T. Mutation of the Carboxy-Terminal Processing Protease in Acinetobacter baumannii Affects Motility, Leads to Loss of Membrane Integrity, and Reduces Virulence. Pathogens 9, 322, doi:10.3390/pathogens9050322 (2020).

7 Roy, R., You, R.-I., Chang, C.-H., Yang, C.-Y. & Lin, N.-T. Carboxy-Terminal Processing Protease Controls Production of Outer Membrane Vesicles and Biofilm in Acinetobacter baumannii. Microorganisms 9, 1336, doi:10.3390/microorganisms9061336 (2021).

8 Kohlmann, F. et al. Structural basis of the proteolytic and chaperone activity of Chlamydia trachomatis CT441. J Bacteriol 197, 211–218, doi:10.1128/JB.02140-14 (2015).

9 Chueh, C.-K. et al. Structural Basis for the Differential Regulatory Roles of the PDZ Domain in C-Terminal Processing Proteases. mBio 10, doi:10.1128/mbio.01129-19 (2019).

10 Su, M. Y. et al. Structural basis of adaptor-mediated protein degradation by the tail-specific PDZ-protease Prc. Nat Commun 8, 1516, doi:10.1038/s41467-017-01697-9 (2017).

11 Hsu, H.-C., Wang, M., Kovach, A., Darwin, A. J. & Li, H. Pseudomonas aeruginosa C-Terminal Processing Protease CtpA Assembles into a Hexameric Structure That Requires Activation by a Spiral-Shaped Lipoprotein-Binding Partner. mBio 13, doi:10.1128/mbio.03680-21 (2022).

12 Hsu, H.-C., Wang, M., Kovach, A., Darwin, A. J. & Li, H. P. aeruginosa CtpA protease adopts a novel activation mechanism to initiate the proteolytic process. The EMBO Journal 43, 1634–1652, doi:10.1038/s44318-024-00069-6 (2024).

13 Lai, T. F., Ford, R. M. & Huwiler, S. G. Advances in cellular and molecular predatory biology of Bdellovibrio bacteriovorus six decades after discovery. Frontiers in Microbiology 14, doi:10.3389/fmicb.2023.1168709 (2023).

14 Thomashow, L. S. & Rittenberg, S. C. WAVEFORM ANALYSIS AND STRUCTURE OF FLAGELLA AND BASAL COMPLEXES FROM BDELLOVIBRIO- BACTERIOVORUS-109J. Journal of Bacteriology 163, 1038–1046 (1985).

15 Lambert, C. et al. Characterizing the flagellar filament and the role of motility in bacterial prey-penetration by Bdellovibrio bacteriovorus. Mol Microbiol 60, 274–286, doi:10.1111/j.1365-2958.2006.05081.x (2006).

16 Seidler, R. J., Starr, M.P.,. Structure of the flagellum of Bdellovibrio bacteriovorus. Journal of Bacteriology 95, 1952–1955, 10.1128/jb.95.5.1952-1955.1968 (1968).

17 Iida, Y. et al. Roles of multiple flagellins in flagellar formation and flagellar growth post bdelloplast lysis in Bdellovibrio bacteriovorus. J Mol Biol 394, 1011–1021, doi:10.1016/j.jmb.2009.10.003 (2009).

18 Morehouse, K. A., Hobley, L., Capeness, M. & Sockett, R. E. Three motAB Stator Gene Products in Bdellovibrio bacteriovorus Contribute to Motility of a Single Flagellum during Predatory and Prey-Independent Growth. Journal of Bacteriology 193, 932–943, doi:10.1128/jb.00941-10 (2011).

19 Shilo, M. & Bruff, B. Lysis of Gram-Negative Bacteria by Host-Independent Ectoparasitic Bdellovibrio bacteriovorus Isolates. Journal of General Microbiology 40, 317–328, doi:10.1099/00221287-40-3-317 (1965).

20 Kaplan, M. et al. Bdellovibrio predation cycle characterized at nanometre-scale resolution with cryo-electron tomography. Nature Microbiology 8, 1267–1279, doi:10.1038/s41564-023-01401-2 (2023).

21 Guerra, J. V. D. S. et al. pyKVFinder: an efficient and integrable Python package for biomolecular cavity detection and characterization in data science. BMC Bioinformatics 22, doi:10.1186/s12859-021-04519-4 (2021).

22 Liao, D. I., Qian, J., Chisholm, D. A., Jordan, D. B. & Diner, B. A. Crystal structures of the photosystem II D1 C-terminal processing protease. Nat. Struct. Biol. 7, 749–753, doi:10.1038/78973 (2000).

23 Zheng, H. et al. CheckMyMetal: a macromolecular metal-binding validation tool. Acta Crystallogr D Struct Biol 73, 223–233, doi:10.1107/S2059798317001061 (2017).

24 Searle, B. C. Scaffold: a bioinformatic tool for validating MS/MS-based proteomic studies. Proteomics 10, 1265–1269, doi:10.1002/pmic.200900437 (2010).

25 Tremblay, C. Y., Vass, R. H., Vachet, R. W. & Chien, P. The Cleavage Profile of Protein Substrates by ClpXP Reveals Deliberate Starts and Pauses. Biochemistry 59, 4294–4301, doi:10.1021/acs.biochem.0c00553 (2020).

26 Fenton, A. K., Kanna, M., Woods, R. D., Aizawa, S. I. & Sockett, R. E. Shadowing the Actions of a Predator: Backlit Fluorescent Microscopy Reveals Synchronous Nonbinary Septation of Predatory Bdellovibrio inside Prey and Exit through Discrete Bdelloplast Pores. Journal of Bacteriology 192, 6329–6335, doi:10.1128/jb.00914-10 (2010).

27 Zhang, H., Zhou, C., Mohammad, Z. & Zhao, J. Structural basis of human 20S proteasome biogenesis. Nat Commun 15, 8184, doi:10.1038/s41467-024-52513-0 (2024).

28 Mabanglo, M. F. & Houry, W. A. Recent structural insights into the mechanism of ClpP protease regulation by AAA+ chaperones and small molecules. J Biol Chem 298, 101781, doi:10.1016/j.jbc.2022.101781 (2022).

29 Duman, R. E. & Lowe, J. Crystal structures of Bacillus subtilis Lon protease. J Mol Biol 401, 653–670, doi:10.1016/j.jmb.2010.06.030 (2010).

30 Clausen, T., Kaiser, M., Huber, R. & Ehrmann, M. HTRA proteases: regulated proteolysis in protein quality control. Nat Rev Mol Cell Biol 12, 152–162, doi:10.1038/nrm3065 (2011).

31 Yonekura, K., Maki-Yonekura, S. & Namba, K. Complete atomic model of the bacterial flagellar filament by electron cryomicroscopy. Nature 424, 643–650, 10.1038/nature01830 (2003).

32 Einenkel, R. et al. The structure of the complete extracellular bacterial flagellum reveals the mechanism of flagellin incorporation. Nat Microbiol 10, 1741–1757, doi:10.1038/s41564-025-02037-0 (2025).

33 Ferenc Vonderviszt, Kanto, S. &, S.-I. A., Keiichi Namba. Terminal regions of flagellin are disordered in solution. Journal of Molecular Biology 209, 127–133, doi:10.1016/0022-2836(89)90176-9 (1989).

34 Altegoer, F. et al. FliS/flagellin/FliW heterotrimer couples type III secretion and flagellin homeostasis. Sci Rep 8, 11552, doi:10.1038/s41598-018-29884-8 (2018).

35 Bhaskar Paidimuddala, J. C., Liman Zhang. Structural basis for flagellin-induced NAIP5 activation. Science Advances 9, doi:10.1126/sciadv.adi8539 (2023).

36 GrüNenfelder, B. R. et al. Identification of the Protease and the Turnover Signal Responsible for Cell Cycle-Dependent Degradation of the *Caulobacter* FliF Motor Protein. Journal of Bacteriology 186, 4960–4971, doi:10.1128/jb.186.15.4960-4971.2004 (2004).

37 Jenal U, S. L. Cell cycle-controlled proteolysis of a flagellar motor protein that is asymmetrically distributed in the Caulobacter predivisional cell. EMBO J 15, 2393–2406 (1996).

38 Ferreira, J. L. et al. γ-proteobacteria eject their polar flagella under nutrient depletion, retaining flagellar motor relic structures. PLoS. Biol. 17, e3000165, doi:10.1371/journal.pbio.3000165 (2019).

39 Hayashi, F., Smith, K., Ozinsky, A. et al. The innate immune response to bacterial flagellin is mediated by Toll-like receptor 5. Nature 410, 1099–1103, 10.1038/35074106 (2001).

40 Sung-il Yoon, O. K., Venkatesh Natarajan, Minsun Hong, Andrei V Gudkov, Andrei L Osterman, Ian A Wilson. Structural basis of TLR5-flagellin recognition and signaling. Science 335, 859–864, doi:10.1126/science.1215584. (2012).

41 Skerker, J. M., Prasol, M. S., Perchuk, B. S., Biondi, E. G. & Laub, M. T. Two-component signal transduction pathways regulating growth and cell cycle progression in a bacterium: a system-level analysis. PLoS Biol 3, e334, doi:10.1371/journal.pbio.0030334 (2005).

42 Lambert, C. & Sockett, R. E. Laboratory Maintenance of Bdellovibrio. Current Protocols in Microbiology 9, 7B.2.1–7B.2.13, 10.1002/9780471729259.mc07b02s9 (2008).

43 Lambert, C. & Sockett, R. E. Nucleases in Bdellovibrio bacteriovorus contribute towards efficient self-biofilm formation and eradication of preformed prey biofilms. FEMS Microbiol Lett 340, 109–116, doi:10.1111/1574-6968.12075 (2013).

44 Fenton, A. K., Lambert, C., Wagstaff, P. C. & Sockett, R. E. Manipulating Each MreB of Bdellovibrio bacteriovorus Gives Diverse Morphological and Predatory Phenotypes. Journal of Bacteriology 192, 1299–1311, doi:10.1128/jb.01157-09 (2010).

45 Lambert, C., Chang, C. Y., Capeness, M. J. & Sockett, R. E. The first bite--profiling the predatosome in the bacterial pathogen Bdellovibrio. PLoS One 5, e8599, doi:10.1371/journal.pone.0008599 (2010).

46 Ducret, A., Quardokus, E. M. & Brun, Y. V. MicrobeJ, a tool for high throughput bacterial cell detection and quantitative analysis. Nat Microbiol 1, 16077, doi:10.1038/nmicrobiol.2016.77 (2016).

47 Remy, O. et al. An optimized workflow to measure bacterial predation in microplates. STAR Protoc 3, 101104, doi:10.1016/j.xpro.2021.101104 (2022).

48 Stewart, P. D. S., Kolek, S. A., Briggs, R. A., Chayen, N. E. & Baldock, P. F. M. Random Microseeding: A Theoretical and Practical Exploration of Seed Stability and Seeding Techniques for Successful Protein Crystallization. Crystal Growth & Design 11, 3432–3441, doi:10.1021/cg2001442 (2011).

49 Stein, N. CHAINSAW: a program for mutating pdb files used as templates in molecular replacement. Journal of Applied Crystallography 41, 641–643, doi:10.1107/s0021889808006985 (2008).

50 Hu, Q. et al. The Orbitrap: a new mass spectrometer. Journal of Mass Spectrometry 40, 430–443, doi:10.1002/jms.856 (2005).

51 Schneider, C. A., Rasband, W. S. & Eliceiri, K. W. NIH Image to ImageJ: 25 years of image analysis. Nature Methods 9, 671–675, doi:10.1038/nmeth.2089 (2012).

52 Crooks, G. E., Hon, G., Chandonia, J. M. & Brenner, S. E. WebLogo: a sequence logo generator. Genome Res 14, 1188–1190, doi:10.1101/gr.849004 (2004).

53 Robert, X. & Gouet, P. Deciphering key features in protein structures with the new ENDscript server. Nucleic Acids Research 42, W320–W324, doi:10.1093/nar/gku316 (2014).

