## Supplementary Information for "A C-terminal Processing Protease Implicated in Flagellin Turnover and Developmental Progression in a Bacterial Predator"

### Supplementary Information Figures

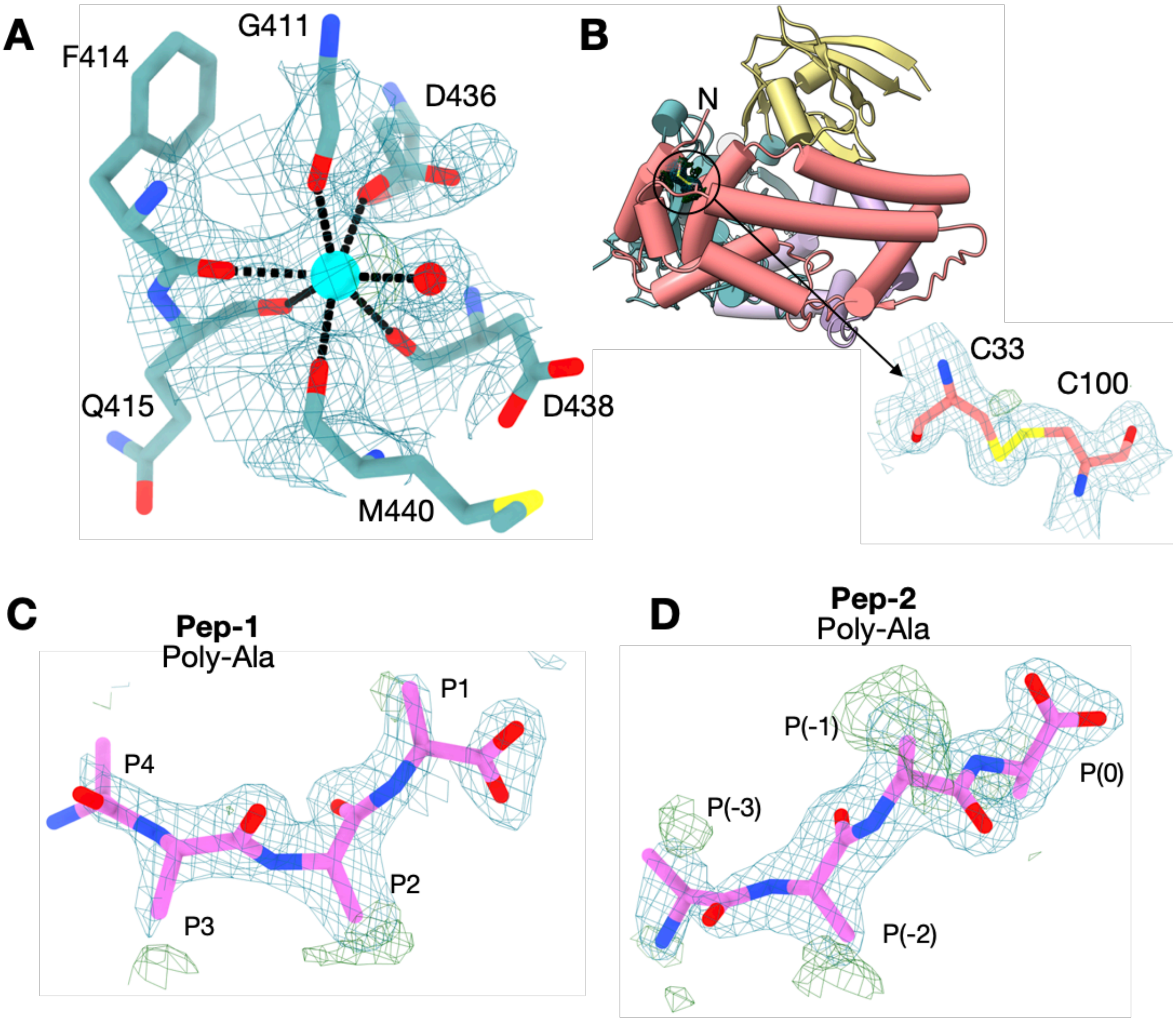

**SI Figure 1. Electron Density Maps**

**(A)** Cation binding site. The $Ca^{2+}$ ion, cyan sphere, is pentagonal bipyramidal coordinated by six hydroxyl and carbonyl groups from surrounding residues, and by one bound water molecule, red sphere. **(B)** The Disulphide bond between residues C33 and C100, located in the N-terminal domain, adds stability to the N-terminus. **(C)** Electron density for the Pep-1 co-purified peptide, which binds in the protease peptide tunnel. **(D**) Electron density for the Pep-2 co-purified peptide, which binds to the PDZ site. Bothe Pep1 and Pep-2 have been modelled as poly-alanine chains, due to the ambiguous electron density maps at the acquired resolution. All electron density maps are illustrate with the 2Fo-Fc map in blue (1.5 σ), with the Fo-Fc map in green (2.5 σ).

**A**

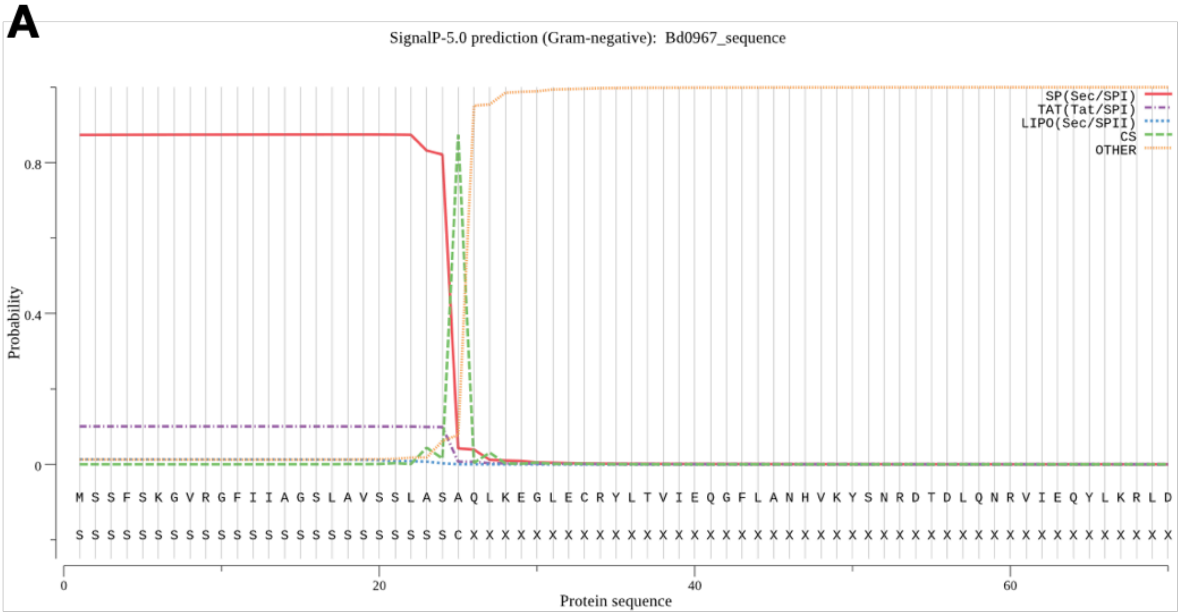

**B** MGSSHHHHHHSSGLVPRGSHQLKEGLECRYLTVIEQGFLANHVKYSNRDTDLQNRVIEQYLKRLDPSKIYLTQGD
VDAIRKSAGNVFDKTKNRDCSFLDAAQKIVLERVKDRSEFAKKYLGKDFKFESSTEFTFDPEKKTWPKDSADANE
YLKKYIQFQIGNYMATDMKMDEAKKNVIKNYERAVKRTADTTQDDLFSGYLDSFARALDPHSSFFSRDVLEDFEIQ
MRLSLEGIGATLSSQDGFTVVEQLVPGGAAAKSGLIEPQDKIVAVGQEKGAMENVIDMDLKDVVKKIRGNKGTKV
RLTILRKSGEGKKRFDVTLTREKVNLEDEAASIIYQDREINGQKKKIGVINFPSFYADSRRGGRSSAADMKKLIKEA
NEKKVDGLVLDLSNNGGGSLEDAVKIAGLFFQTGNVVKQSSKNEGRAESALRDTDPMVDWSGPLVVLTSRISAS
ASEIVSGTLQDYKRAVVVGGDHTYGKGSVQSVLPIPNNLGAIKVTVGMFFVPGGKSTQHRGVDADIVLPGPFSAD
DIGEKYMDYSLPPKTIESFLSPDAYVKEGPGAWKEIKPEWLKSLRERSGERVAKNDEFKKIVEELNKAKARGKVIR
VSEVLKDKNEKEKKDKAKKTASKAKKNEEYLKRPDIMEAENVLLDLIQLEDGKSLVPQQKQANAK

**Key**
Thrombin cleavable His-6 tag
Bd0967 sequence

**C** MGSSHHHHHHSSGLVPRGSHMGMRISTNVSAINAQRTMVNSQREIGKSMSQLASGSRINKAADDAAGLAISENL
KSQIRSLGQASRNANDGISMVQTAEGGLSEISNILTRMRELGVQASSDTIGDTERGFLDKEVQQLKSEAQRITQTT
RFGTTKLLDGSGDSFDFQVGINNDPEADRISFNAGETNASTSSLGIDGFDFSSKTGAQDALAAIDTAQSQVNGYR
ANLGALQNRLQSTVDNLGVQHENISAANSRIRDTDVAAATAETTRNQVLLQANTSVLSQANAMPNSALRLIG

**Key**
Thrombin cleavable His-6 tag
Bd0606 sequence
RLIG sequence truncated in ΔLIG and ΔRLIG

#### SI Figure 2. Bd0967 signal peptide prediction and construct design

**(A)** Signal IP 5.0 graphical output, illustrating the predicted signal peptide with cleavage site at residue A24. **(B)** The Bd0967 expression construct comprised residues 25–673, excluding the predicted signal peptide, and was cloned downstream of an N-terminal His$_6$-tag followed by a thrombin cleavage site. **(C)** Bd0606 was cloned and expressed as a full-length construct comprising residues 1-277, fused to an N-terminal thrombin cleavable His-6 tag. ΔRLIG and ΔLIG C-terminal truncation mutants are also shown.

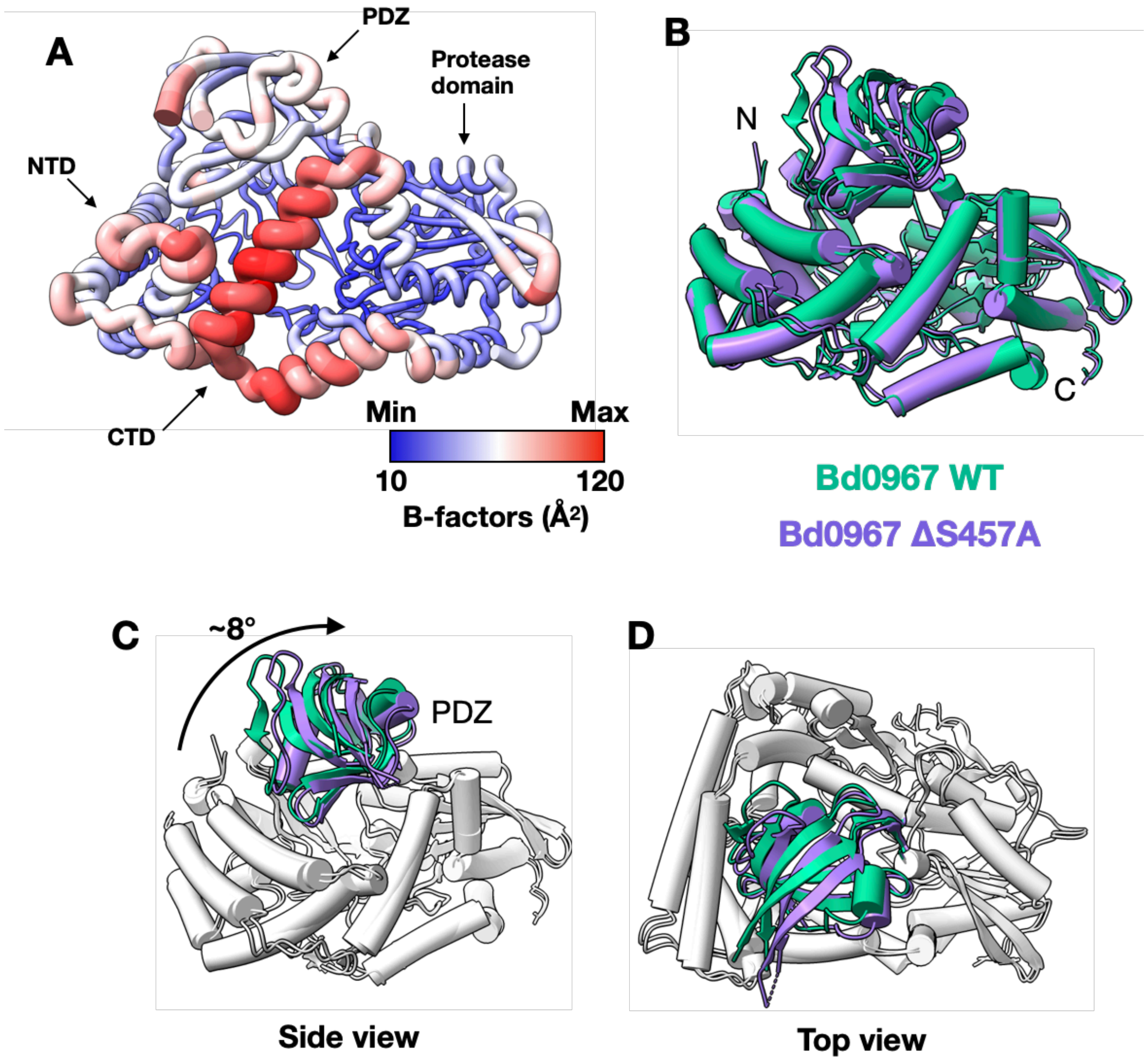

**SI Figure 3. Comparison of Bd0967 structures**

**(A)** B-factors shown in worm form on the structure of Bd0967. Higher B-factors are associated with the distal regions of the NTD and CTD, as well as the PDZ domain. **(B)** Structural alignment of Bd0967 WT at 2.04 Å  and Bd0967$_{\Delta S457A}$ at 2.58 Å, by least-squares structural superposition (LSQ) about the protease domain (residues 333-566). WT is coloured green and S457A mutant is coloured purple. **(C)** Illustration of the modest PDZ domain conformational change, which rotates at ~8° about the hinge region axis (at its greatest). The colour depiction of the PDZ domain is the same as for A, but the rest of the structure has been coloured white to highlight the movement**.** **(D)** Showing PDZ flexibility from a top down view.

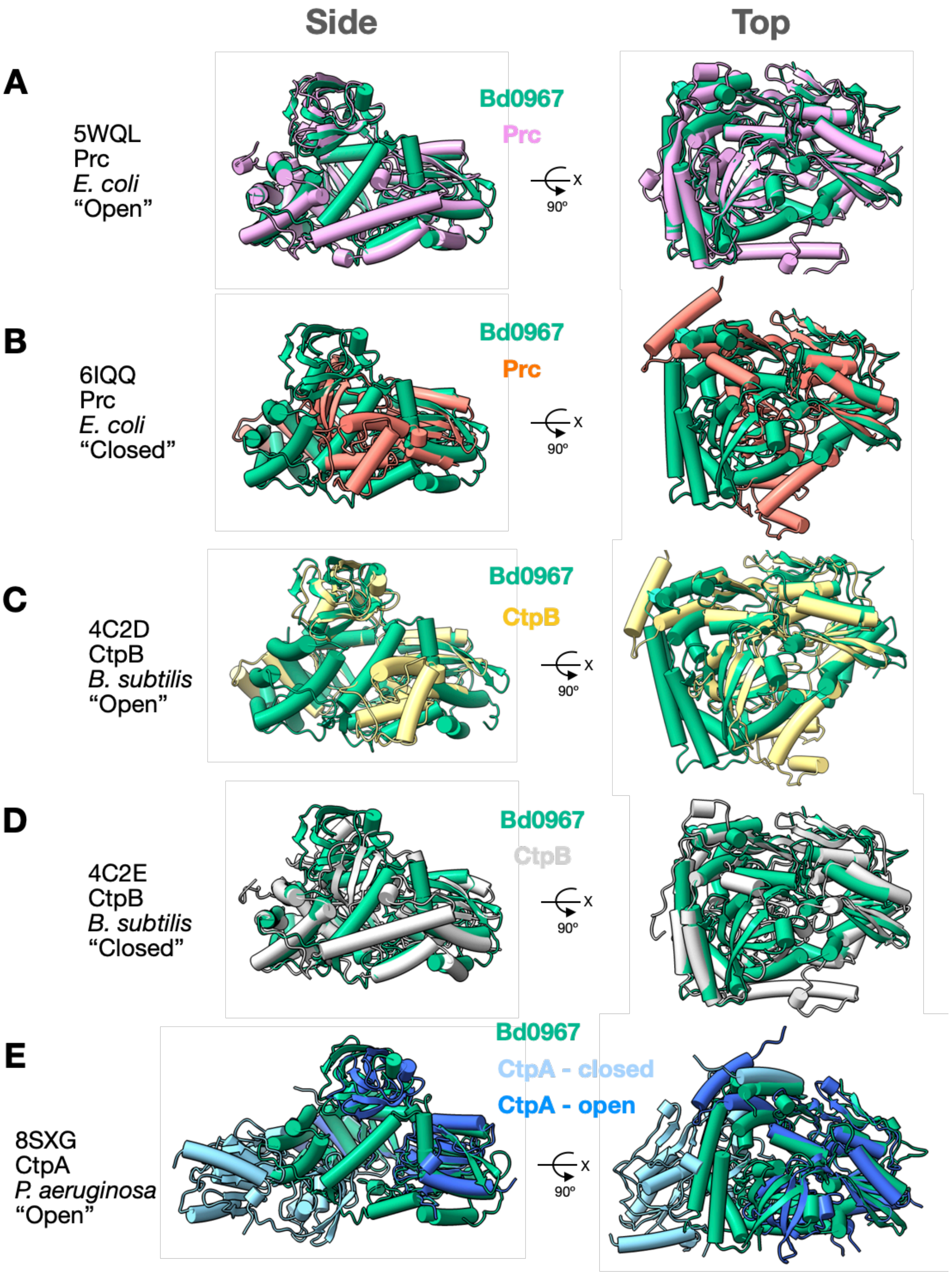

**SI Figure 4. Bd0967 structure comparison to CTP family structures**

Structural alignment of Bd0967 (green) to CTP-family proteases using least-squares structural superposition. **(A)** Prc from *Escherichia coli* in the open form; **(B)** Prc from E.coli in the closed form; **(C)** CtpB from *Bacillus subtilis* in the open form; (D) CtpB in the closed form. **(E)** CtpA from *Pseudomonas aeruginosa* in the open form in chain B, chain A PDZ domain is in a closed form.

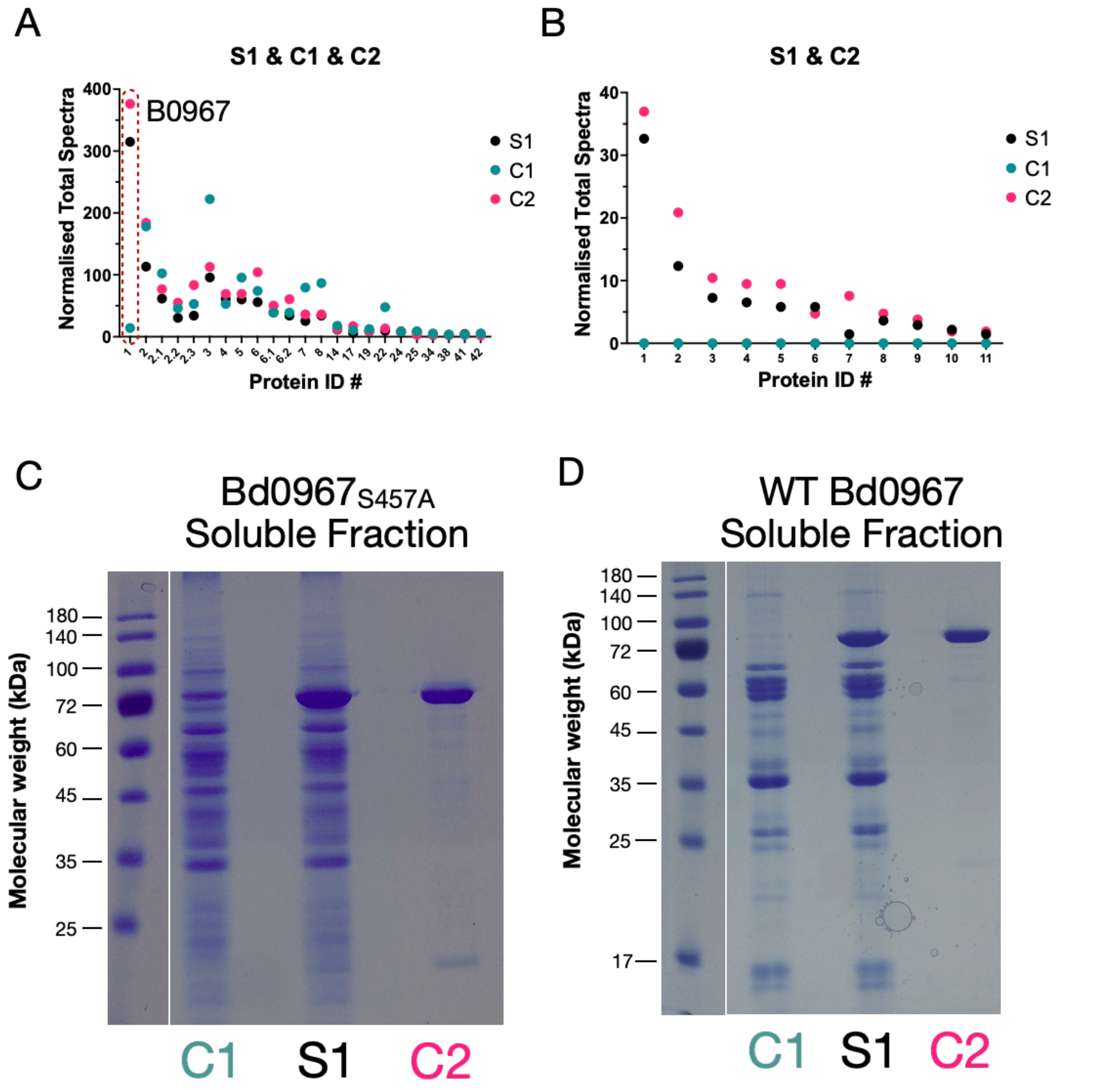

**SI Figure 5. Additional MS data plots and SDS-PAGE**

**(A)** Graph showing the normalised total spectra, for samples identified in S1 & C1 & C2. Bd0967 (highlighted by the red dashed shape) was very highly enriched in the S1 & C2 samples, and very low abundance in the C1 sample, as expected. **(B)** Graph showing the normalised total spectra, for samples identified in S1 & C2. These samples were all identified as contaminating *E. coli* species. **(C)** SDS-PAGE of pulldown conducted with Bd0967$_{S457A}$ and soluble fraction. No unique bands in the S1 lane compared to C1 and C2. **(D)** SDS-PAGE of pulldown conducted with WT Bd0967 and soluble fraction. No unique bands in the S1 lane compared to C1 and C2 lanes.

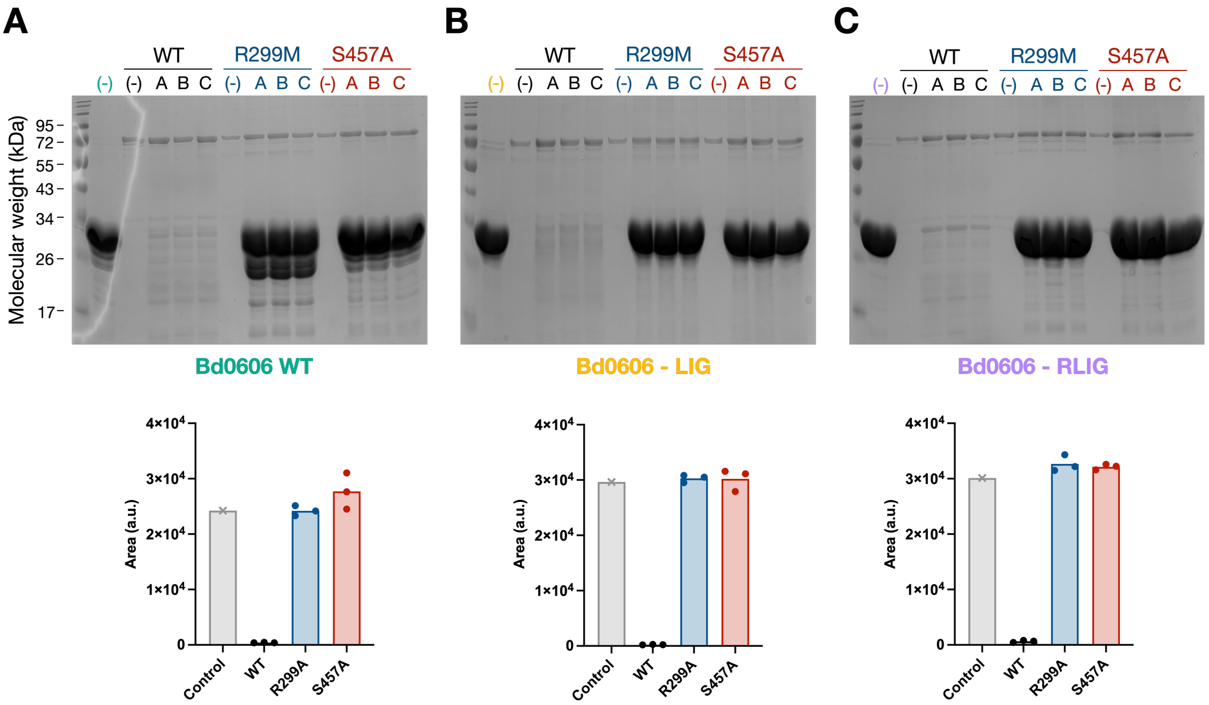

#### SI Figure 6. Mutational analysis of key residues

SDS-PAGE gel images are shown for each Bd0606 variant, with corresponding band intensity quantification displayed below. Each gel contains three independent replicate reactions performed in parallel and resolved on the same SDS–PAGE gel. **(A)** Bd0606 degradation by WT Bd0967, Bd0967$_{R299M}$ and Bd0967$_{S457A}$. Nucleophile substituted Bd0967$_{S457}$ is inactive, whilst Bd0967$_{R299M}$ has very low activity compared to WT. The appearance of a lower molecular weight intermediate suggests limited activity. **(B)** Bd0606-LIG degradation by WT Bd0967, Bd0967$_{R299M}$ and Bd0967S457A. Bd0967$_{R299M}$ has no activity compared to WT. **(C)** Bd0606-LIG degradation by WT Bd0967, BD0967$_{R299M}$ and Bd0967S457A. Bd0967$_{R299M}$ has no activity compared to WT. Bd0967$_{R299M}$ shows minimal activity to WT Bd0606 substrate but not for C-terminal truncation mutants, suggesting R199 residue is important for substrate recognition particularly in less efficient substrates.

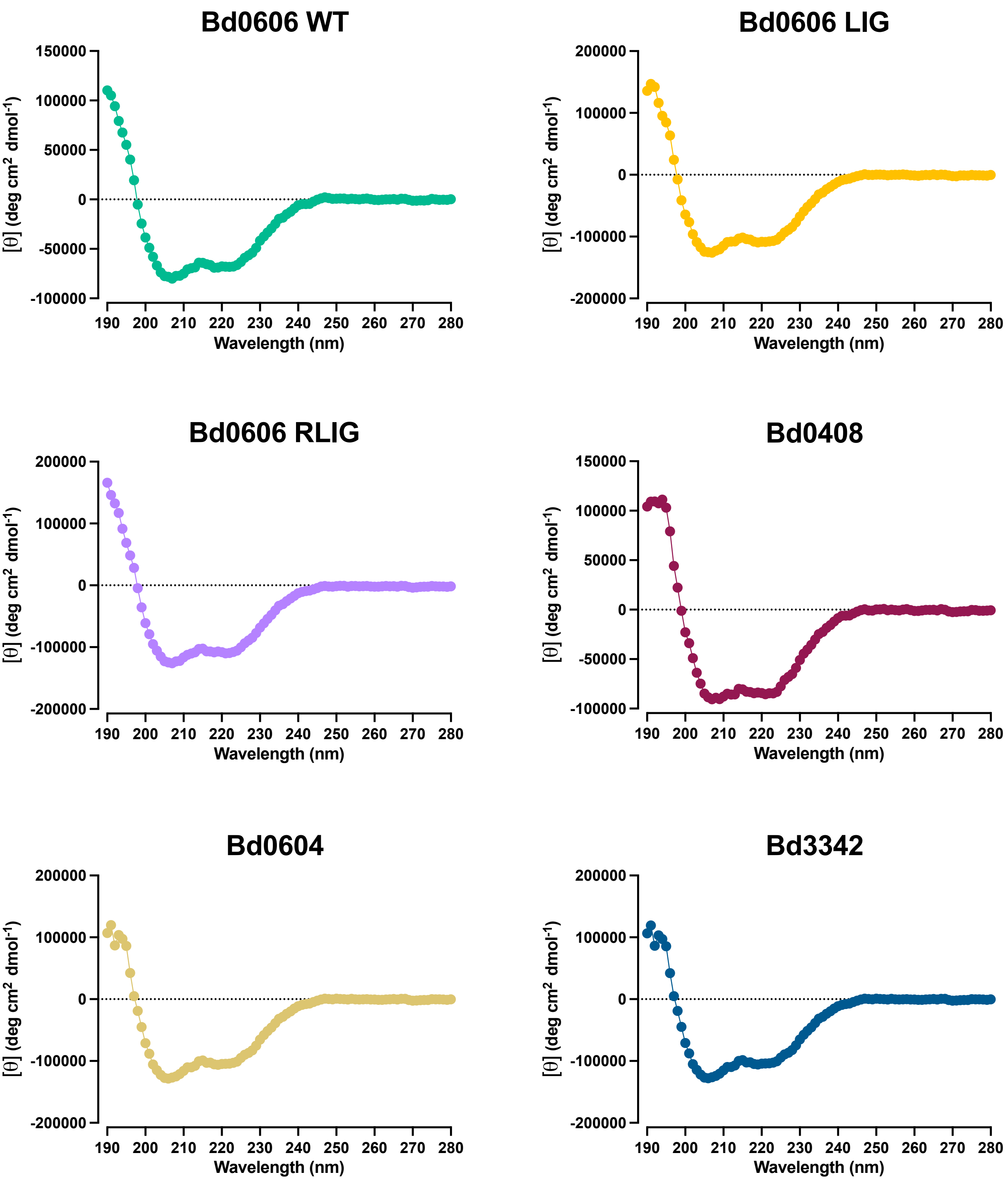

**SI Figure 7. Far-UV circular dichroism spectra of recombinant flagellin proteins.**

Spectra were recorded from 260–190 nm in 20 mM sodium phosphate buffer (pH 8.0) using a 1 mm quartz cuvette. Each spectrum represents the average of 10 scans and is shown as mean residue ellipticity ([θ], deg·cm$^2$·dmol$^{-1}$) following buffer subtraction. All proteins display characteristic minima at ~208 and ~222 nm, consistent with predominantly α-helical secondary structure expected of folded flagellins.

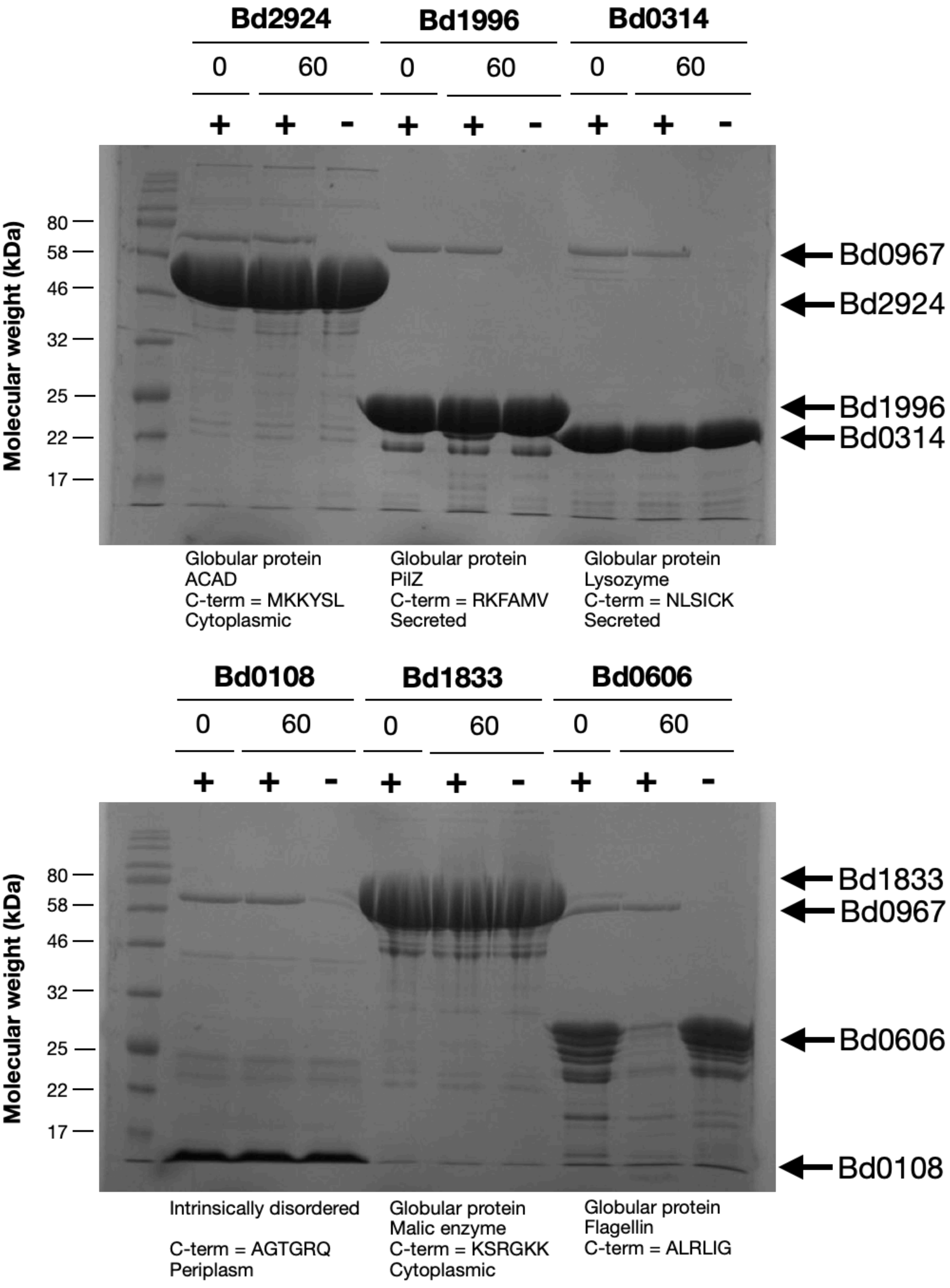

**SI Figure 8. Bd0967 substrate specificity**

SDS-PAGE gel images showing the activity of Bd0967 towards a selection of recombinantly expressed *Bdellovibrio* proteins. The c-terminal sequence of each protein is presented alongside their predicted cellular location, structural fold and protein identity. The proteolytic assay was performed under standard assay conditions, alongside a negative control (no Bd0967 enzyme). Samples were taken at 0 and 60 minutes. The only protein which showed degradation over the 60 minute period was Bd0606.

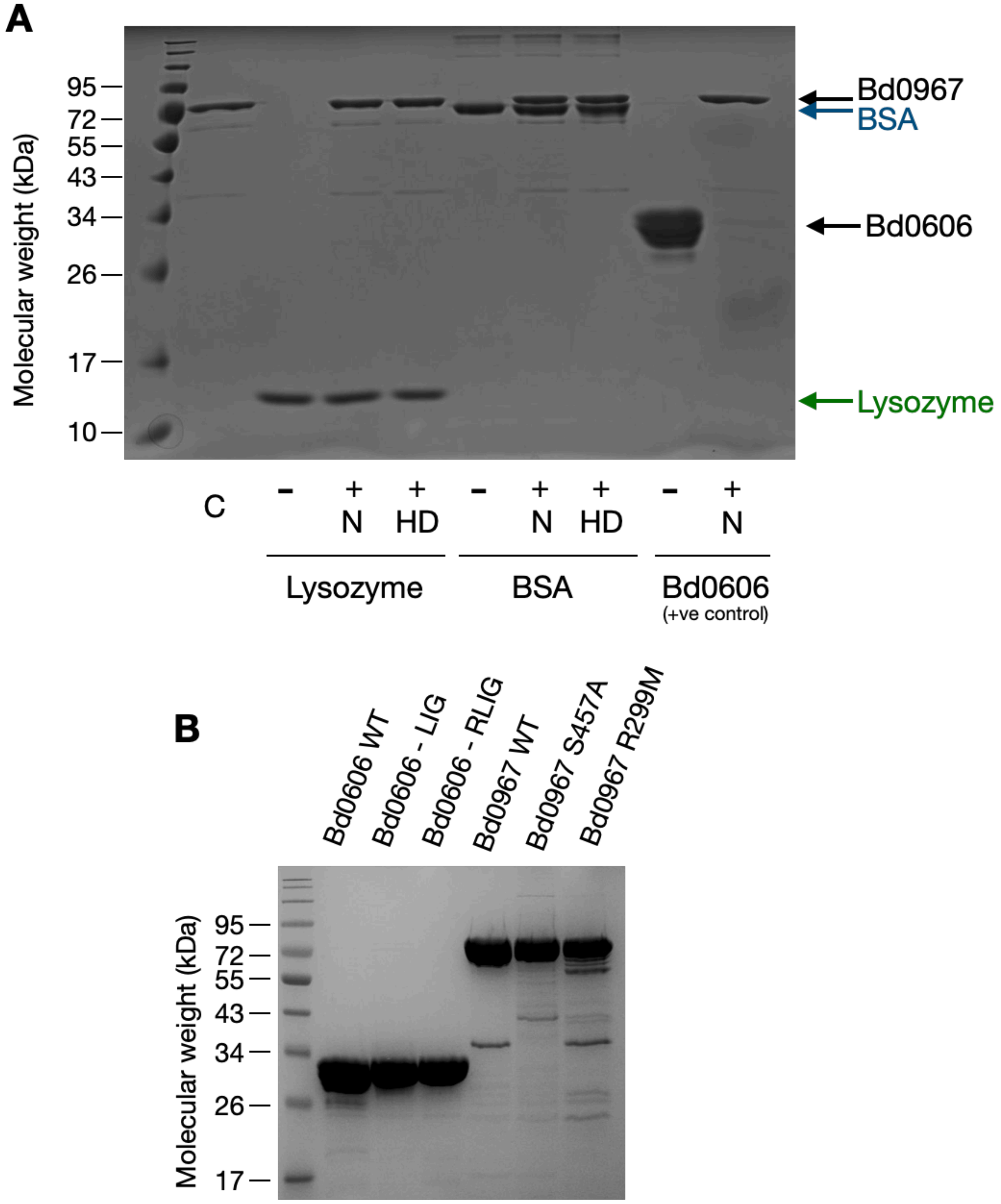

**SI Figure 9. Controls and Sample purity**

**(A)** SDS-PAGE showing the activity of Bd0967 towards native (N) and heat-denatured (HD) proteins, under standard assay conditions at 37 ºC for 60 minutes. A Bd0967 only control (C), was run along a positive control, Bd0606, which was completely degraded within the assay period. No degradation of Lysozyme or BSA in their native or heat denatured states was detected, demonstrating these are not substrates of Bd0967. This suggests Bd0967 selects substrates base on structure and sequence rather than unfolded state. **(B)** SDS-PAGE gel showing the purity of proteins used in this investigation. Proteins loaded at a concentration of 50 μM (10 μl loaded).

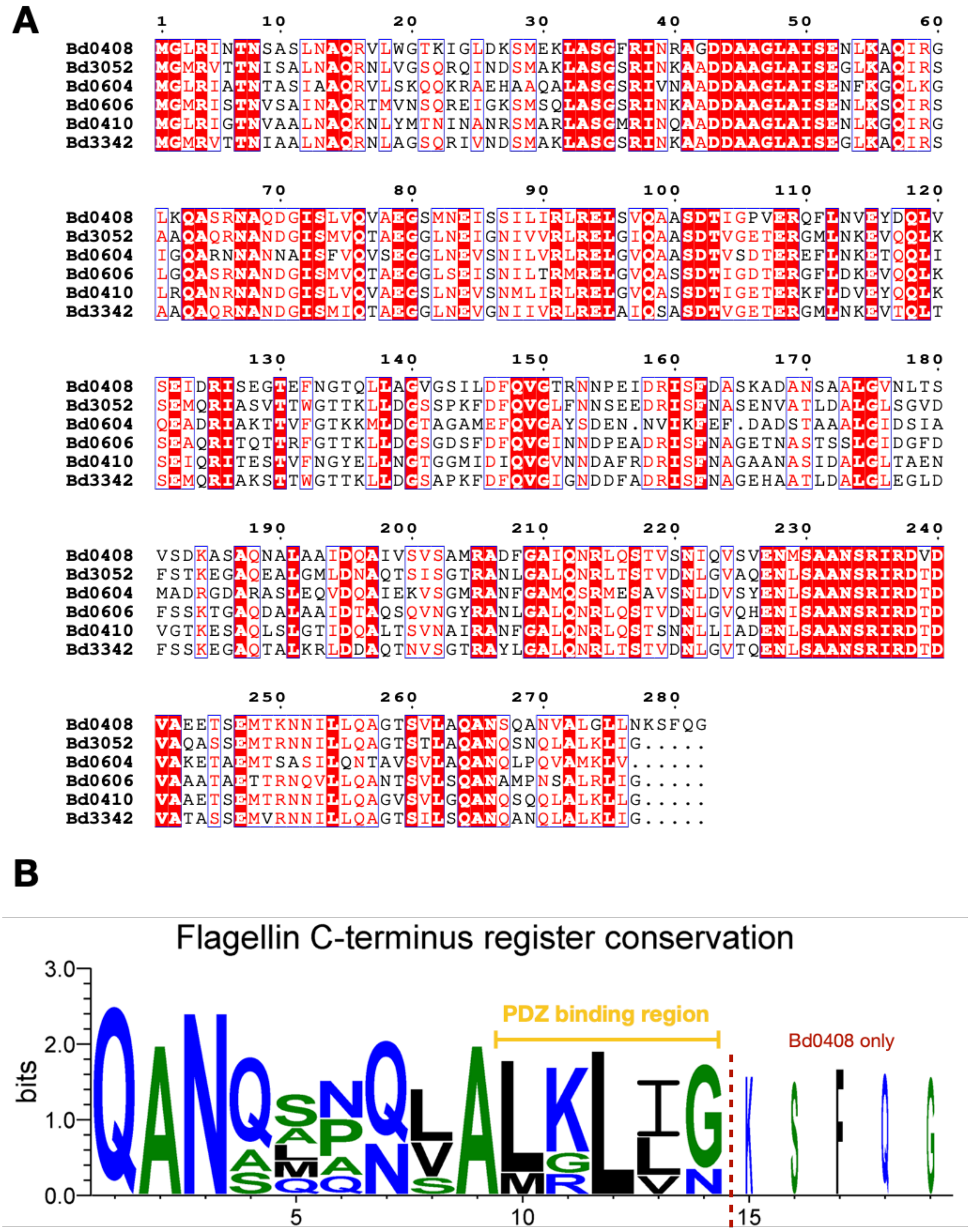

**SI Figure 10. Flagellin sequence alignment**

**(A)** The six *Bdellovibrio bacteriovorus* flagellins were aligned using Clustal omega multiple sequence alignment tool. The flagellins are organised from proximal (Bd0408) to distal (Bd3342). The C-terminus aligns well, apart from Bd0408, which has an additional 5 residues at the C-terminus. **(B)** An illustration of the conservation of the C-terminus, created using WebLogo3. This shows clear conservation of residues at the C-terminus, with the exception of Bd0408. The flagellin C-terminal motif is represented by the region highlighted as the PDZ binding region.

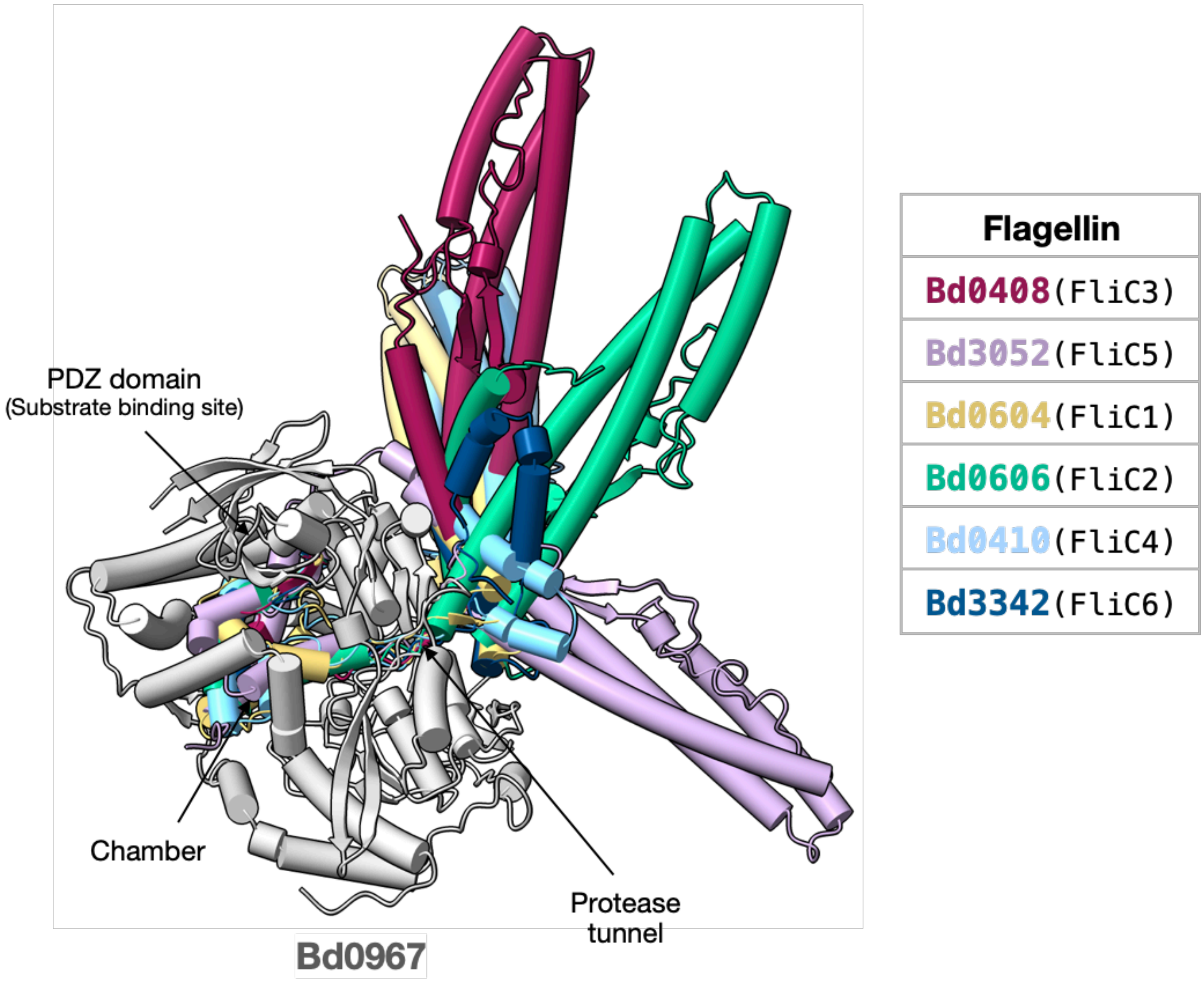

**SI Figure 11. AlphaFold3 model of Bd0967 interaction with Bdellovibrio flagellins**

Alphafold 3 was used to predict and model the interaction between Bd0967 and the Bdellovibrio flagellins. Each flagellin is predicted to interact with Bd0967 in a comparable way. The C-terminus interacts directly with the PDZ domain substrate binding site. A region of approximately 35 amino acids is predicted to pass through the protease tunnel into the the cavity of Bd0967. The remainder of the flagellins alpha helical structure is modelled in several different conformations, suggesting prediction confidence of this region compared to the C-terminus.

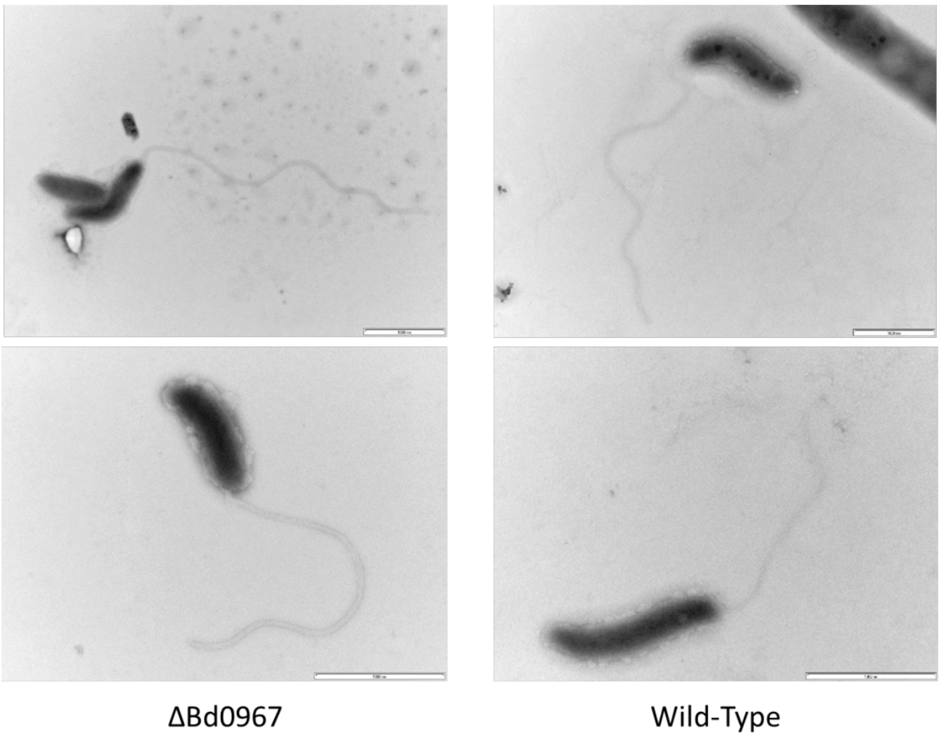

**SI Figure 12. Electron micrographs – strain similarity**

Electron micrographs demonstrating the majority of ΔBd0967 cells were morphologically similar to wild-type HD100. Cells were stained with 0.5% uranyl acetate. Images are representative of three independent experiments. Scale bars are 1 µm.

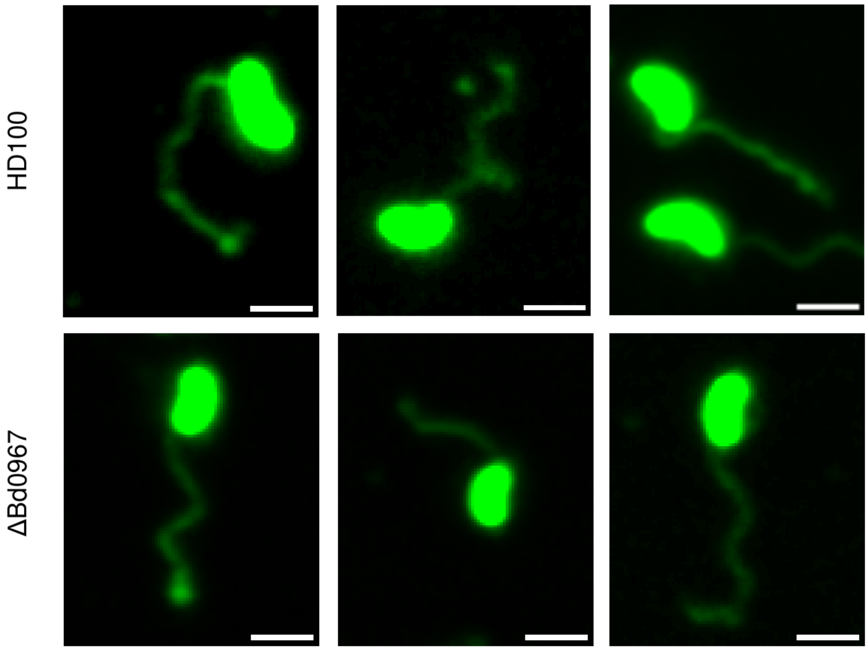

**SI Figure 13. Epifluorescent micrographs – strain similarity**

Epifluorescent micrographs of attack phase *B. bacteriovorus* HD100 wild- type and Δ*Bd0967* mutant cells stained with Vibrant Dio membrane stain (false-colored green) demonstrating the majority of Δ*Bd0967* cells were morphologically similar to wild-type HD100. Scale bar = 1μm.

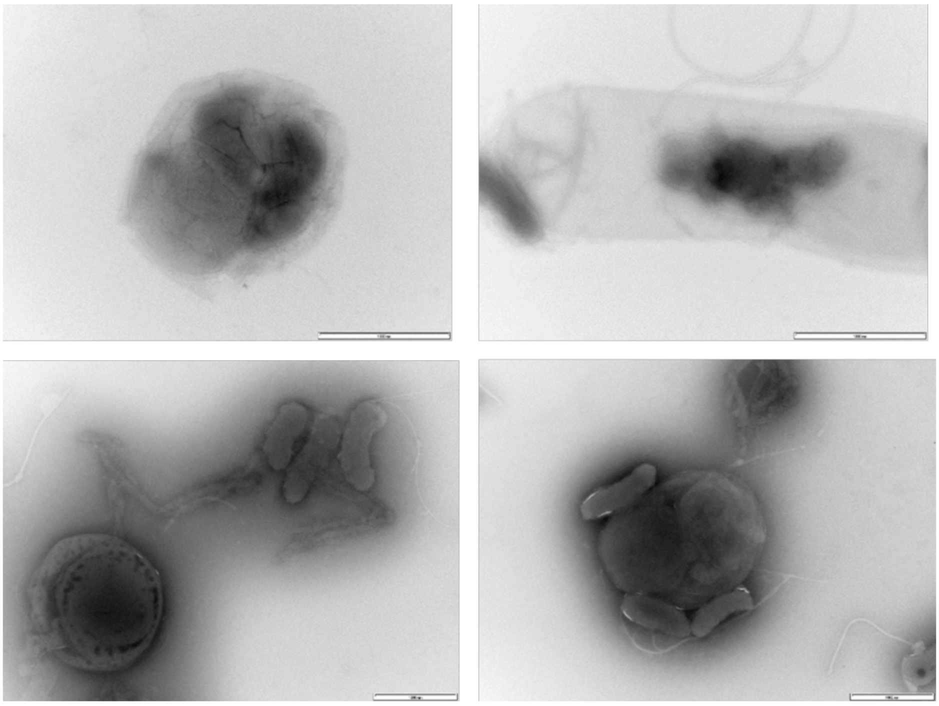

**SI Figure 14. Electron micrographs – stalled bdelloplasts**

Electron micrographs showing stalled bdelloplasts with apparently deformed Δ*Bd0967* cells within. Cells were stained with 0.5% uranyl acetate. Images are representative of three independent experiments. Scale bars are 1 μm.

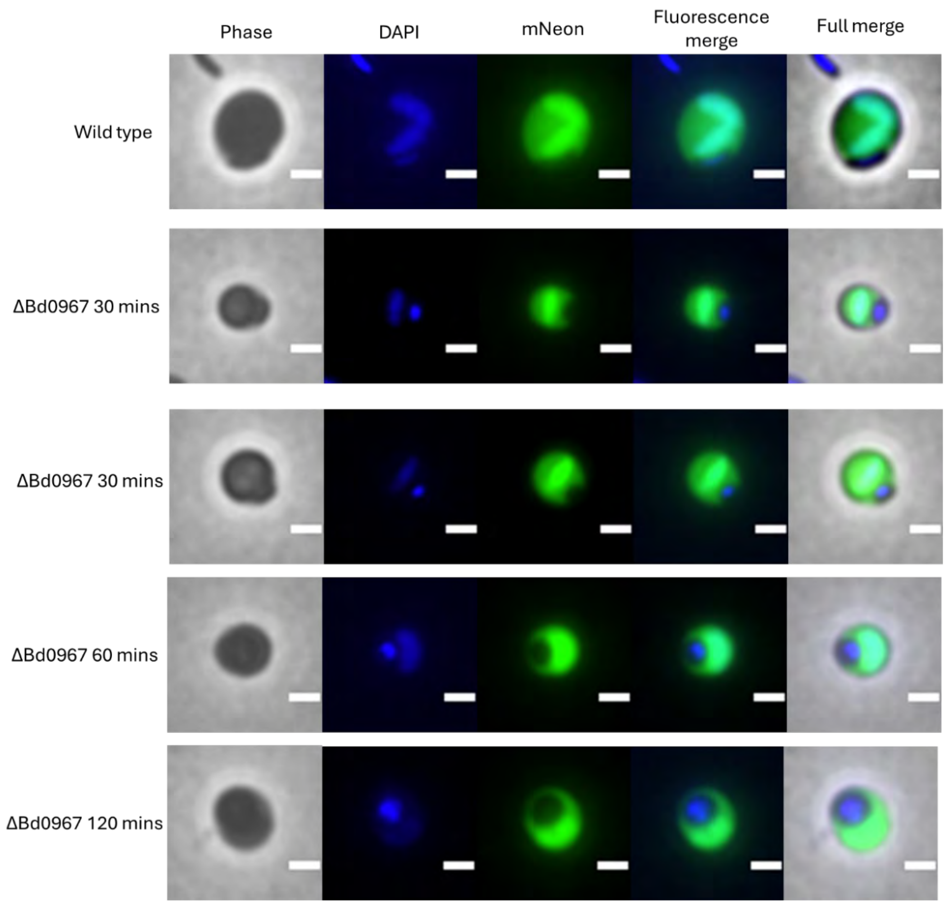

**SI Figure 15. Epifluorescent micrographs – aberrant growth in bdelloplasts**

Epiflourescent and phase contrast micrographs of wild-type bdelloplasts and aberrant growth of Δ*Bd0967* within bdelloplasts at 30-120 minutes post-mixing of *Bdellovibrio* and prey *E. coli*. Prey periplasms are labelled with pMal::mNeonGreen and DNA is stained with DAPI (blue). Images are representative of three independent experiments. Scale bars are 1 µm

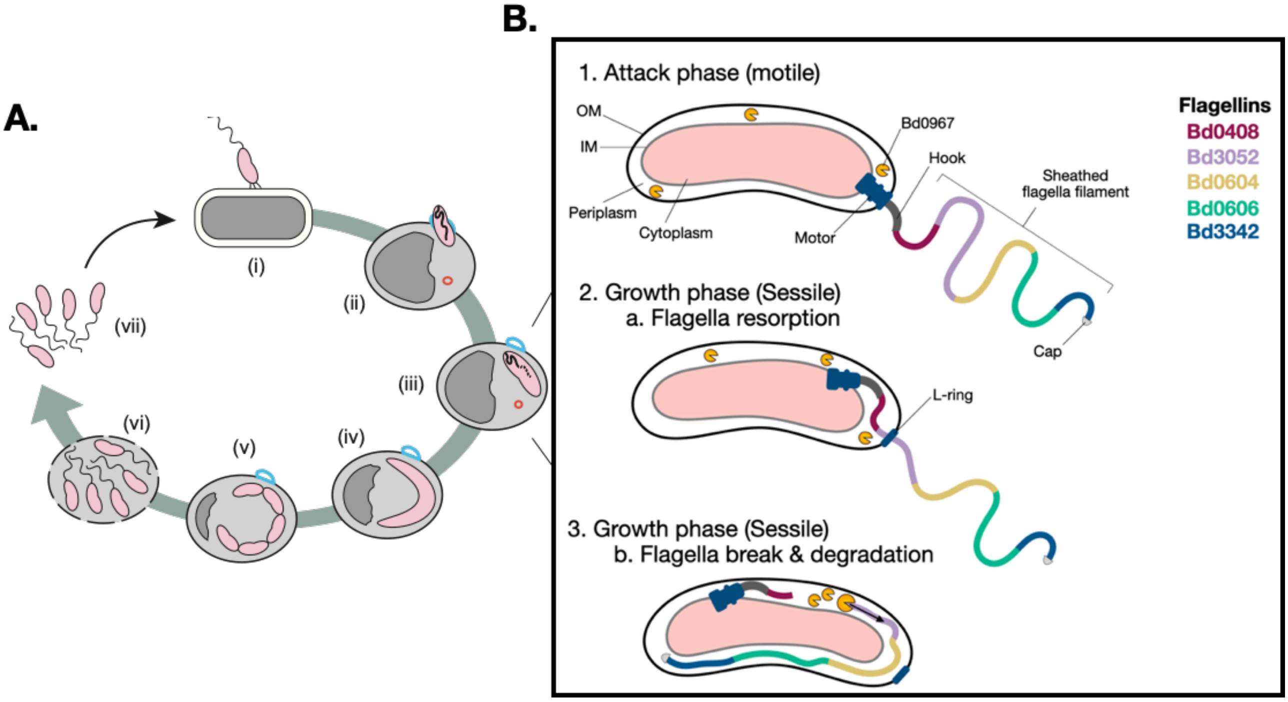

**SI Figure 16. Proposed model for Bd0967-mediated flagellin degradation, after flagellar resorption, during the predatory lifecycle of Bdellovibrio bacteriovorus.**

**A)** Schematic overview of the predatory lifecycle of *B. bacteriovorus*. Following attachment to a Gram-negative prey cell (i), the predator invades the prey periplasm and forms a bdelloplast (ii). Intraperiplasmic growth, chromosome replication, and filament elongation occur within the bdelloplast (iii–iv), followed by septation and progeny maturation (v). The prey cell is subsequently lysed (vi), releasing motile progeny that re-enter the attack phase (vii). *B. bacteriovorus* cells are shown in pink and prey cells in grey, with the prey cytoplasm and periplasm indicated in dark and light grey, respectively. **(B)** Model for flagellar resorption and degradation by Bd0967 during the transition from attack phase to intraperiplasmic growth. In attack-phase cells, Bd0967 is distributed throughout the periplasm. Upon prey invasion, previous cryo-electron tomography studies have shown that the flagellar filament is resorbed into the predator cell envelope[1]. During this process, the flagellar motor disengages from the basal body and migrates within the inner membrane, while the L-ring remains associated with the outer membrane. Retraction of the flagellar sheath into the outer membrane is proposed to facilitate movement of the filament into the periplasm (2a). Subsequent filament destabilisation exposes the N- and C-terminal regions of flagellin subunits at the proximal end. Bd0967 recognises accessible C-terminal motifs and initiates proteolysis of the filament, consistent with the observed localisation of Bd0967-mCherry to discrete periplasmic foci during growth. Progressive degradation of flagellin subunits is proposed to expose additional C-termini, enabling sequential depolymerisation and turnover of the resorbed filament.

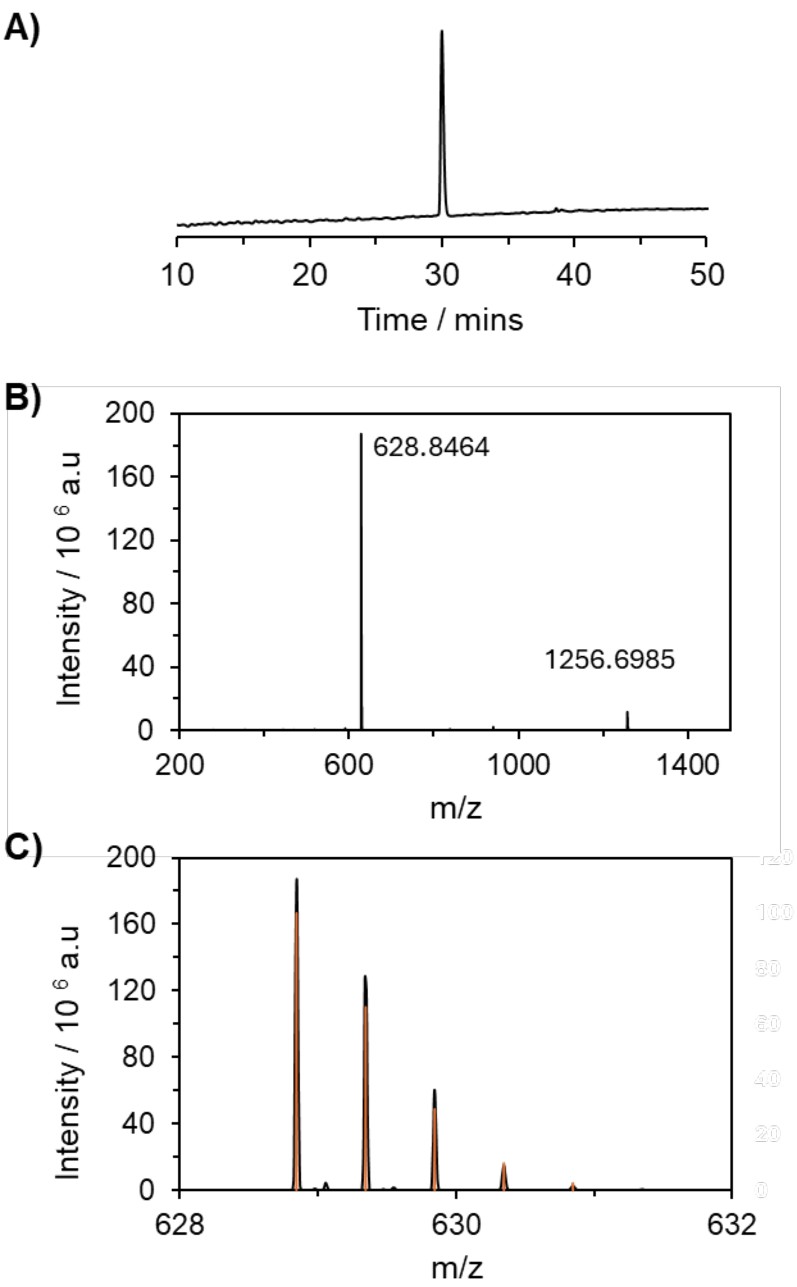

**SI Figure 17. Peptide Synthesis – Inhibitor A (NAMPNSALRLIG)**

**A)** Analytical C18 reversed-phase HPLC trace of purified H-NAMPNSALRLIG-OH using a linear 0 – 60% MeCN + 0.05% TFA in $H_2O$ + 0.05% TFA gradient over 60 minutes and monitored at 210 nm. Electrospray ionization mass spectra, of **B)** the charge envelope and **C)** the $[M+2H]^{2+}$ isotopic distribution of purified H-NAMPNSALRLIG-OH with the experimental (black) and theoretical (orange) spectra shown. The average mass of the peptide is 1256.5 Da.

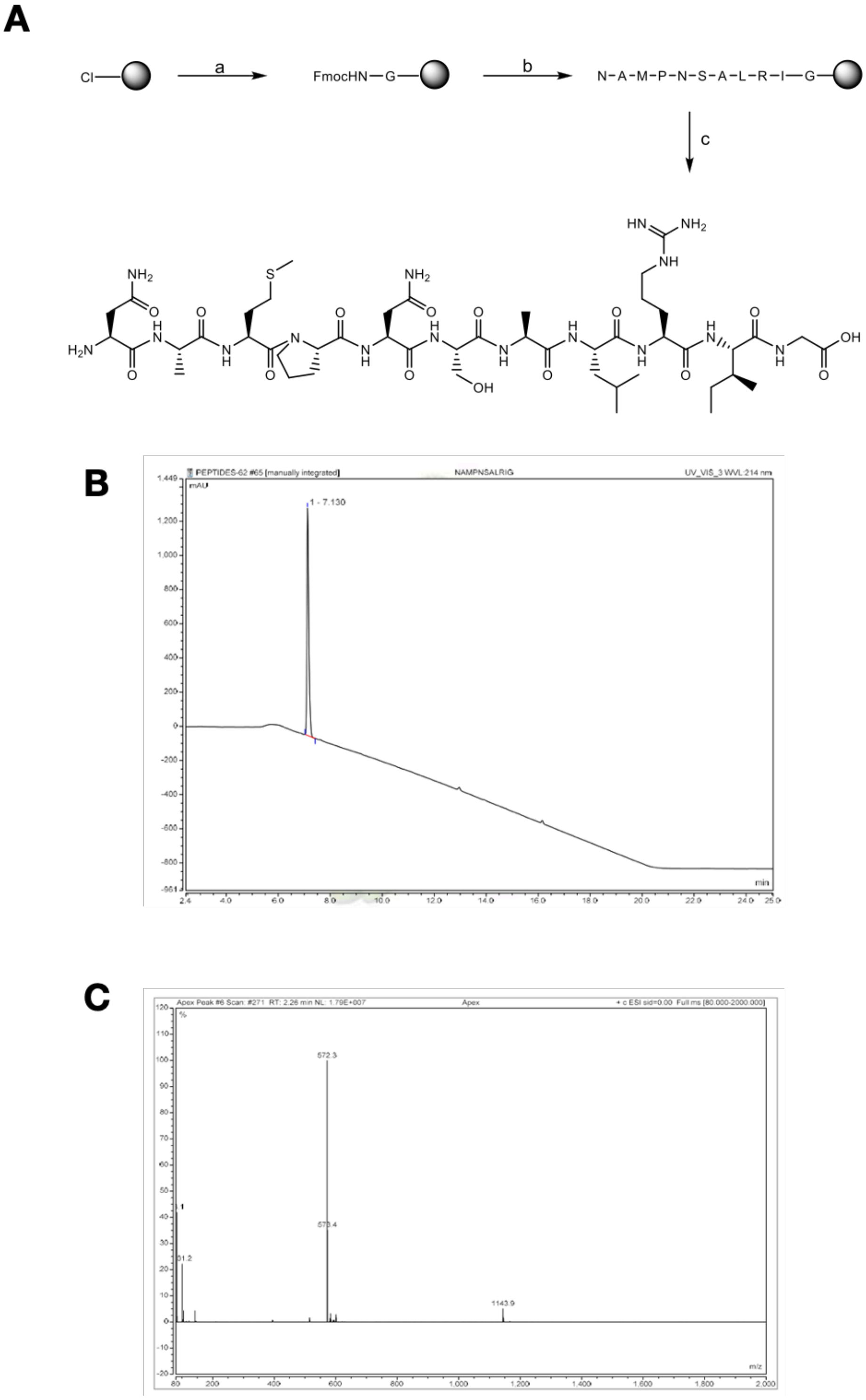

**SI Figure 18. Peptide Synthesis – Inhibitor B (NAMPNSALRIG)**

**A)** Schematic overview of the workflow used for the synthesis of NAMPNSALRIG. **B)** HPLC trace of HPLC purified NAMPNSALRIG (gradient: 5-95% ACN in 25 min using A: 0.1% HCOOH in water, B: MeCN). **C)** MS spectra from of HPLC purified NAMPNSALRIG. Exact Mass calcd. for $C_{47}H_{82}N_{16}O_{15}S$ = 1142.5866, found $M+H^+$ = 1143.9 and $M/2 + H^+$ = 572.3

### Supplementary Information Tables

#### Supplementary Table 1. Primers

| | Primer | Sequence (5' to 3') | Purpose |
|---|---|---|---|
| **Strain Construction** | 967KOupF | CGTTGTAAAACGACGGCCAGTGCCAAGGG ACAATTGGAGTTAGG | Deletion of *bd0967* |
| | 967KOupR | CCTCAAATCACAGCTGATCCTTCCATGGCT C | Deletion of *bd0967* |
| | 967KOdownF | TGGAAGGATCAGCTGTGATTTGAGGAAAC GTTTGTTG | Deletion of *bd0967* |
| | 967KOdownF | GGAAACAGCTATGACCATGATTACGAGCCC AAGCACCATGACG | Deletion of *bd0967* |
| | pK18mobsac F | AGAGGATCCCCGGGTACCG | *bd0967-mCherry* fusion |
| | pK18mobsac R | AGAGTCGACCTGCAGGCATG | *bd0967-mCherry* fusion |
| | pk18mobsac-upBd0967-F | TCGGTACCCGGGGATCCTCTAAGGATAGAA ACTTCCACCTTGGCG | *bd0967-mCherry* fusion |
| | pk18mobsac-upBd0967-R | CACCATGAGCTCGAGGATGTCCTTGGCGT TGGCCTGTTTCTG | *bd0967-mCherry* fusion |
| | lnk-mcherry-Bd0967-F | CCAAGGACATCCTCGAGCTCATGGTGAGC AAGGGCGAGG | *bd0967-mCherry* fusion |
| | lnk-mcherry-downBd0967-R | CAAACGTTTCCTCAAATCACACTACTTGTA CAGCTCGTCCATGCC | *bd0967-mCherry* fusion |
| | mcherry-downBd0967-F | CATGGACGAGCTGTACAAGTAGTGTGATTT GAGGAAACGTTTGTTGTTGTTG | *bd0967-mCherry* fusion |
| | downBd0967-pk18mobsac-R | CATGCCTGCAGGTCGACTCTCAGCCCAAG CACCATGACG | *bd0967-mCherry* fusion |
| **Recombinant Expression Constructs** | Bd0967_Full_Fwd | GCAGCGGCCTGGTGCCGCGCGGCAGCCA TCAACTGAAAGAAGGCCTGGAGTGC | Full length Bd0967 construct in pET28a |
| | Bd0967_Full_Rev | CTCAGTGGTGGTGGTGGTGGTGCTCGAG CTACTTGGCGTTGGCCTGTTTCTG | Full length Bd0967 construct in pET28a |
| | Bd0606_Fwd | GCAGCGGCCTGGTGCCGCGCGGCAGCCA TATGGGAATGAGAATTTCTACG | Full length Bd0606 construct in pET28a |
| | Bd0606_Rev | GCACTAAGACTGATTGGTTAACTCGAGCAC CACCACCACCACCACTGAG | Full length Bd0606 construct in pET28a |
| **Mutagenesis** | Bd0967_S457A_Fwd | TATCTCTGCTGCCGCTTCCGA | Site directed mutation of S457 to alanine |
| | Bd0967_S457A_Rev | CGGCTGGTCAATACCACC | Site directed mutation of S457 to alanine |
| | Bd0967_R299M_Fwd | CAAAAAAATCATGGGTAACAAGGGCACGAA AGTG | Site directed mutation of R299 to methionine |
| | Bd0967_R299M_Rev | ACAACGTCTTTCAAATCC | Site directed mutation of R299 to methionine |

#### Supplementary Table 2. Plasmids

| Plasmid | Description | Source |
|---|---|---|
| pK18*mobsacB* | Suicide vector (kanR, *lac*Za, *sacB*) used for crossovers into the *B. bacteriovorus* genome | Schafer *et al.* 1994[2] |
| pdeltaBd0967 | Upstream and downstream fragments around *bd0967*gene to generate unmarked gene deletion | This study |
| pcompBd0967 | *bd0967*gene with upstream and downstream fragments to return the gene for complementation | This study |
| pAKF220-mNeon | Backlighting *E. coli* cells with mNeon green fluorescence. | Makowski et al., 2019[3] |
| pK18*mobsacB-bd0967-mcherry* | bd0967-linker-mCherry translational fusion | This study |
| pET28a | Recombinant gene expression<br>N-term thrombin cleavable site | Lab stock |

#### Supplementary Table 3. Strains

| Strains | Description | Source |
|---|---|---|
| *E. coli* NEBa (DH5a) | *E. coli* cloning strain ( *fhuA2Δ(argF-lacZ)U169 phoA glnV44 Φ80Δ(lacZ)M15 gyrA96 recA1 relA1* ) | New England Biolabs (C2987) |
| *E. coli* BL21 (DE3) | *E. coli* expression strain *(fhuA2 [lon] ompT gal (λ DE3) [dcm] ∆hsdS*<br>*λ DE3 = λ sBamHIo ∆EcoRI-B int::(lacI::PlacUV5::T7 gene1) i21 ∆nin5)* | New England Biolabs (C2527H) |
| *E. coli* SHuffle® T7 | *E. coli* expression strain ( F´ *lac, pro, lacI*$^{q}$ */ Δ(ara-leu)7697 araD139 fhuA2 lacZ::T7 gene1 Δ(phoA)PvuII phoR ahpC* galE (or U) galK λatt::pNEB3-r1-cDsbC* (Spec$^{R}$, *lacI*$^{q}$) *ΔtrxB rpsL150*(Str$^{R}$) *Δgor Δ(malF)3 )* | New England Biolabs (C3026J) |
| *E. coli* S17-1 | *E. coli* strain (thi, pro, hsdR-, hsdM+, recA; integrated plasmid RP4- Tc::Mu-Kn::tn) | Hanahan D., 1983[4] |
| *B. bacteriovorus* HD100 | *B. bacteriovorus* Type strain, genome-sequenced, wild-type | Rendulic S, *et al.*, 2004[5] |
| *B. bacteriovorus* HD100 deltaBd0967 | *B. bacteriovorus* HD100 with a markerless deletion of *bd0967* | This study |
| *B. bacteriovorus* HD100 compBd0967 | *B. bacteriovorus* HD100 deltaBd0967 complemented by returning *bd0967* to its original locus | This study |
| *E. coli* S17-1 mNeon | *E. coli* S17-1 containing plasmid pAKF220-mNeon | Makowski et al., 2019[3] |

| | | |
|---|---|---|
| *B. bacteriovorus* HD100 *Bd0967*-mCherry | *B. bacteriovorus* HD100 with a *bd0967-mCherry* C-terminal fusion at native locus, merodiploid | This study |
| *E.coli* DH5α mTurquoise2 | *E. coli* DH5α containing plasmid pFED343-*Ptac-mTurquoise2* | Mukherjee lab |

#### Supplementary Table 4. Crystallisation conditions

| | **Condition** |
|---|---|
| Bd0967 (seed stock) | Morpheus G4 -<br>0.1 M Carboxylic acids, 0.1 M Buffer System 1 pH 6.5, 37.5 % v/v Precipitant Mix 4 |
| Bd0967 | Morpheus G4 -<br>0.1 M Carboxylic acids, 0.1 M Buffer System 1 pH 6.5, 37.5 % v/v Precipitant Mix 4 |
| Bd0967 S457A | Morpheus G12 -<br>0.1 M Carboxylic acid mix, 0.1 M Buffer system 3 pH 8.5, 37.5 % v/v Precipitant mix 4 |

**Supplementary Table 5. Crystallographic Data Table**

| | | Bd0967 | Bd0967S457A |
|---|---|---|---|
| **Accession code** | | 30XH | 30XJ |
| **Data Collection** | | | |
| | Resolution (Å) | 106.26 - 2.04 (2.07 - 2.04) | 38.42 - 2.58 (2.62 - 2.58) |
| | Space group | P 1 | P 1 |
| | Cell Dimensions a, b, c (Å) | 73.42, 95.75, 114.11 | 72.88, 95.02, 113.26 |
| | α, β, γ (°) | 73.48, 71.30, 81.89 | 77.68, 71.80, 81.94 |
| | Total reflections | 848316 (18620) | 304174 (15674) |
| | Unique reflections | 179707 (8159) | 87121 (4331) |
| | Multiplicity | 4.7 (2.3) | 3.5 (3.6) |
| | Completeness (%) | 97.8 (89.6) | 98.5 (98.1) |
| | Mean I/sigma(I) | 4.8 (1.2) | 7.5 (1.1) |
| | R-meas | 0.184 (0.453) | 0.134 (1.254) |
| | R-pim | 0.079 (0.28) | 0.071 (0.654) |
| | CC1/2 | 0.952 (0.733) | 0.995 (0.448) |
| **Refinement** | | | |
| | R-free | 0.238 | 0.252 |
| | R-work | 0.196 | 0.212 |
| | Total non-hydrogen atoms | 21763 | 20122 |
| | Total macromolecule atoms | 19934 | 19921 |
| | Total ligand atoms (Ca) | 4 | 4 |
| | Total solvent atoms | 1825 | 197 |
| | Protein molecules per ASU | 4 | 4 |
| | Residues per protein | 637 | 637 |
| | RMS(bonds) (Å) | 0.008 | 0.01 |
| | RMS(angles) (°) | 1.2 | 1.64 |
| | Ramachandran favoured (%) | 97.49 | 96.98 |
| | Ramachandran allowed (%) | 2.39 | 2.86 |
| | Ramachandran outliers (%) | 0.12 | 0.16 |
| | Average B-factor (Å2) | 44.19 | 63.64 |

*Values in parentheses are for the high-resolution shell
**R-value test set size =  5

#### Supplementary Table 6. Fitted data

| Substrate | Model | Y0 (AU) | X0 (min) | K (min-1) | 95% CI | $T_{1/2}$ (min) | 95% CI | $R^2$ |
|---|---|---|---|---|---|---|---|---|
| **Bd0606 WT** | One phase decay | 27464 | | 0.0520 | 0.0454 - 0.0596 | 13.3 | 11.6 - 15.3 | 0.9883 |
| **Bd0606 LIG** | Plateau followed by one phase decay | 24803 | 10.0 | 0.0307 | 0.0198 - 0.0418 | 22.6 | 16.6 - 35.1 | 0.9802 |
| **Bd0606 RLIG** | Plateau followed by one phase decay | 22944 | 13.9 | 0.0348 | 0.0321 - 0.0448 | 19.9 | 15.5 - 21.6 | 0.9899 |

#### Supplementary Table 4. Flagellin construct

### Supplementary methods section

#### Supplementary Methods 1 – Peptide Synthesis of Inhibitor A (NAMPNSALRLIG)

Peptide Synthesis and Purification of H-NAMPNSALRLIG-OH: Fmoc protected amino acids were purchased from Sigma Aldrich, AGTC Bioproducts, Cambridge Reagents or Novabiochem. The peptide was synthesized using an automated microwave assisted CEM Liberty Blue on Wang resin (1.51 mmol/g substitution, 0.25 mmol scale). The synthesis was carried out using standard Fmoc-amino acid protocols[6] where deprotection conditions (20% piperidine in DMF) with 0.1 M 1-hydroxybenzotriazole in the deprotection mixture and standard O-(benzotriazol-1-yl)-N,N,N',N'-tetramethyluronium hexafluorophosphate/1-hydroxybenzotriazole coupling conditions were used[7]. The peptide was cleaved as previously reported[2] and purified using a C18 preparative HPLC column using a linear 10 – 60% acetonitrile + 0.05% TFA in $H_2O$ + 0.05% gradient.

**Characterization of H-NAMPNSALRLIG-OH:**
The peptide was characterized by mass spectrometry using a nanoelectrospray ionisation (TriVersa NanoMate, Advion Interchim Scientific, Ithaca, NY) installed on a Waters Synapt G2-S Mass Spectrometer (Waters Corporation, Wilmslow, UK), operating at a resolution of 20,000 FWHM. Data was collected in positive ion mode.

#### Supplementary Methods 2 – Peptide Synthesis of Inhibitor B (NAMPNSALRIG)

##### I. Materials

All L amino acids, oxyma pure, Diisoproplycarbodiimide (DIC), Thioanisole, Triisopropylsilane (TIS), Trifluoroacetic acid (TFA) and Diisopropylethylamine (DIPEA) were purchased from Fluorochem, UK. Dimmethylformamide (DMF) peptide synthesis grade was purchased from Rathburn chemicals. Diethyl ether, Dichloromethane, Formic acid 98-100% purity, Water (HPLC garde) and Acetonitrile (HPLC grade) were purchased from Fisher Scientific. 2-Chlorotritylchloride resin (manufacturer's loading: 1.20 mmol/g) was purchased from Iris Biotech. All chemicals were used without further purification.

##### II. Equipment used for the analysis and purification of compounds:

All peptides were analysed on a Thermo Scientific Dionex Ultimate 3000 RP-HPLC equipped with a Phenomenex Gemini NX C18 110 Å (150 x 4.6 mm) column using the following buffer systems: A: 0.1% HCOOH in HPLC water. B: MeCN using a flow rate of 1 ml/min. The column was flushed with 95% A for 5 min prior to an injection and was flushed for 5 min with 95% B and 5% A after the run was finished.

Peptides were analysed using the following gradient: 95% A for 2 min. 5-95% B in 25 min. 95% B for 5 min. 5% A for 4 min.

Peptides were purified using the same gradient as mentioned above, on a Biotage® Isolera one flash purification system with a flow rate of 25mL/min.

LC-MS data were collected on a Thermo Scientific Dionex Ultimate 3000 RP-UPLC instrument with a Phenomenex Kinetex C18 100Å column (50 x 2.1 mm, 2.6μm at 30 °C) connected to a Thermo Scientific™ ISQ™ EC Single Quadrupole Mass Spectrometer with a flow rate of 0.6 mL/min with the following solvent systems: (A): 0.1% HCOOH in $H_2O$ and (B) MeCN. The column was flushed with 95% A for 2 min, then a gradient from 5% A to 95% B over 6 min was used, followed by 2 min of flushing with 95% B.

#### III. Synthetic Scheme of peptide sequence NAMPLSALRIG

**Step a -** Commercially available 2-Chlorotrityl chloride resin (manufacturer's loading = 1.2 mmol/g, 1g resin) was swelled in DCM in a reactor. To this resin was added 4 eq. Fmoc-Gly-OH/8 eq. DIPEA in DCM and the reactor was shaken for 3h. The loading was calculated to be 0.6 mmol/g, (830mg resin, 0.5mmol). Any unreacted resin was capped with MeOH:DIPEA:DCM = 1:2:7 by shaking for 1h.

**Step b** - Deprotection cycles were performed 3min at 50°C followed by 10min at RT to remove the Fmoc. All amino acids were coupled using 4 eq. Amino Acid, 4 eq. DIC/Oxyma using a microwave peptide synthesiser. Coupling time was 10 min at 50°C.

**Step c -** The resin and side-chain protecting groups of the peptide were then cleaved off using TFA:Thioanisole:TIS:H2O = 90:5:2.5:2.5 by stirring for 2h. The peptide was precipitated using cold $Et_2O$ (-20°C) and centrifuging at 7800 rpm to obtain a white solid. This solid was further purified using the equipment and methods described in section II and pure fractions were pooled and freeze dried to obtain a white solid 68mg, 12% yield.

### Supplementary References

1 Kaplan, M. *et al.* Bdellovibrio predation cycle characterized at nanometre-scale resolution with cryo-electron tomography. *Nature Microbiology* **8**, 1267-1279, doi:10.1038/s41564-023-01401-2 (2023).

2 Andreas Schäfer, A. T., Wolfgang Jäger, Jörn Kalinowski, Georg Thierbach, Alfred Pühler. Small mobilizable multi-purpose cloning vectors derived from the Escherichia coli plasmids pK18 and pK19: selection of defined deletions in the chromosome of Corynebacterium glutamicum. *GENE* **145**, 69-73, doi:10.1016/0378-1119(94)90324-7 (1994).

3 Makowski, L. *et al.* Dynamics of Chromosome Replication and Its Relationship to Predatory Attack Lifestyles in Bdellovibrio bacteriovorus. *Appl Environ Microbiol* **85**, doi:10.1128/AEM.00730-19 (2019).

4 Hanahan, D. Studies on transformation of Escherichia coli with plasmids. *J MOL BIOL* **166**, 557-580, doi:10.1016/s0022-2836(83)80284-8 (1983).

5 Rendulic, S. *et al.* A predator unmasked: Life cycle of Bdellovibrio bacteriovorus from a genomic perspective. *Science* **303**, 689-692, doi:10.1126/science.1093027 (2004).

6 Chan, W. & White, P. D. *Fmoc Solid-Phase Peptide Synthesis: A Practical Approach*. Vol. 222 (2000).

7 Peacock, A. F. *et al.* Gold-phosphine binding to de novo designed coiled coil peptides. *J Inorg Biochem* **117**, 298-305, doi:10.1016/j.jinorgbio.2012.05.010 (2012).
